# Lineage-selective suicide gene system enables post-engraftment editing of cell therapy composition

**DOI:** 10.64898/2026.06.03.729995

**Authors:** Jennifer Jin, Chiara Pavan, Niamh Moriarty, Dmitry A. Ochinnikov, Gabriella Farrell, Andrew T. Quattrocchi, Cameron PJ Hunt, Clare L Parish

## Abstract

Human pluripotent stem cell (hPSC)-derived therapies are advancing rapidly toward clinical application, yet heterogeneity of transplanted cell populations remains a major barrier to safety, predictability and scalability. Existing strategies to mitigate this risk either incompletely eliminate proliferative cells or ablate the entire graft, thereby compromising therapeutic benefit. Here we present NeuroGuard, a lineage-selective suicide gene platform that decouples safety from efficacy by preserving functional neurons while enabling inducible elimination of all other cell types after transplantation. NeuroGuard integrates an inducible caspase-9 system with NEUROD1-driven Cre recombination, protecting post-mitotic neurons from apoptosis while rendering non-neuronal and proliferative populations susceptible to ablation. *In vitro*, activation of the system enriched neuronal content to >90% and increased dopaminergic neuron proportion >3-fold. Following transplantation of ventral midbrain progenitors, timed activation eliminated proliferative and glial populations, resulting in compact, neuron-enriched grafts without loss of dopaminergic neuron number, target innervation or behavioural recovery in Parkinsonian rodents. Single-cell transcriptomics confirmed selective removal of non-neuronal lineages while preserving neuronal identity and maturation programs. This work establishes a generalizable framework for post-engraftment editing of cell therapy composition, providing a versatile strategy to enhance the safety and functional predictability of regenerative therapies.

## INTRODUCTION

Human pluripotent stem cell (hPSC)-derived cell therapies are rapidly progressing toward clinical translation, offering new opportunities for the treatment of neurological and other degenerative diseases. In Parkinson’s disease (PD), transplantation of ventral midbrain (vm) dopamine progenitors has shown promising outcomes in preclinical models and early-phase clinical trials, with evidence of graft survival, integration and functional benefit ^1–5^. More broadly, advances in stem cell engineering and manufacturing are enabling increasingly sophisticated and programmable cell therapy products across disease areas ^6–8^.

Despite these advances, a central limitation persists: the heterogeneity of cells within these transplants. Current differentiation protocols reliably generate regionally specified progenitors, yet these populations remain intrinsically mixed ^9^. Following transplantation, grafts typically contain a minority of dopaminergic neurons alongside diverse off-target populations, including other neuronal subtypes, glia and proliferative progenitors ^10–14^. This heterogeneity has important implications for both safety and efficacy, with proliferative or off-target/poorly specified cells contributing to graft overgrowth and unpredictable functional outcomes ^15–17^.

Efforts to improve graft purity have largely focused on eliminating unwanted cells and/or highly proliferative cells prior to transplantation using cell sorting approaches. While these methods have enabled enrichment, they remain limited by the absence of a sufficiently restrictive protein/gene to isolate the desired population ^18–22^. Moreover, the risks of proliferative cells evading cell sorting or activation of quiescent stem cells within the graft remain, meaning graft composition cannot be fully controlled prior to implantation.

To address these limitations, inducible suicide genes have emerged as an important safety strategy for engineered cell therapies ^23^. These systems enable pharmacological activation of apoptosis within donor cells and are increasingly used across regenerative medicine and immunotherapy contexts, including adoptive cell therapies where drug-inducible safety switches are essential for managing toxicity and improving clinical control ^7,23^. Among these, inducible caspase-9, cytosine deaminase/5-fluorocytine and herpes simplex virus thymidine kinase are most widely adopted – targeted at killing either all cells within a graft or only those actively proliferating ^23^. Whilst we and others have demonstrated success in improving safety through this approach ^7,24–29^, strategies targeting proliferating cells incompletely resolve graft heterogeneity, while systems that ablate all donor cells eliminate therapeutic benefit ^7,30^.

These limitations reflect a broader challenge in cell therapy, that is the need for strategies enabling precise, programmable control over cell identity and composition *in vivo*, and represents a major focus of next-generation synthetic biology and engineered cell therapy platforms, particularly in the CAR-T therapies ^8,31,32^.

To address this, we engineered NeuroGuard, a lineage-selective suicide gene platform designed to enable post-engraftment editing of graft composition. The system integrates inducible caspase-9 (iCasp9) together with Cre recombinase driven by a neuronal transcription factor (NEUROD1), that is expressed in post-mitotic neurons. Upon tamoxifen-induced Cre activation, the suicide gene is excised specifically in neurons, rendering them resistant to subsequent apoptosis, induced by a chemical dimerizer. By contrast, non-neuronal and proliferative cells retain the suicide gene and remain susceptible to apoptosis. This design enables selective protection of the functional neuronal component of the graft, while eliminating all other cell populations.

We apply NeuroGuard to hPSC-derived vm progenitor grafts as a clinically relevant model of neuronal cell therapy, targeted at PD. We demonstrate efficient lineage-specific protection and selective ablation *in vitro* and show that post-transplant activation yields compact, neuron-enriched grafts without compromising dopaminergic neuron number or behavioural recovery. Single-nuclei transcriptomic profiling further confirms selective elimination of non-neuronal populations within mature grafts. Together, these findings establish NeuroGuard as a broadly applicable strategy for decoupling safety from efficacy and enabling programmable control of cell therapy composition *in vivo*, a central goal in emerging regenerative and synthetic cell therapy technologies.

## RESULTS

### Generation, validation and vm differentiation of the AAVS1-iCasp9:Neurod1-CreERT2 hPSC line

The new hPSC line, engineered to express iCasp9 at the AAVS1 safe harbour locus together with a Cre recombinase cassette at the NEUROD1 locus, was subsequently referred to as NeuroGuard. The transgenic PSC line was generated via a two-phase CRISPR/Cas9 genome engineering strategy. First, an inducible caspase 9 (iCasp9) donor construct with a PGK-driven puromycin resistance cassette was targeted to the *AAVS1* safe harbour locus using dual-nickase CRISPR/Cas9, enabling selection and PCR-based validation of correctly integrated clones (***Figure 1Ai; Figure S1***). In the second phase, a CreERT2–Neomycin donor cassette was inserted at the *NEUROD1* locus in validated *AAVS1*-iCasp9 clones using a similar approach, followed by puromycin (PURO) and G418 selection and PCR screening for correct integration (***Figure 1Aii; Figure S1***.

**Figure 1.**
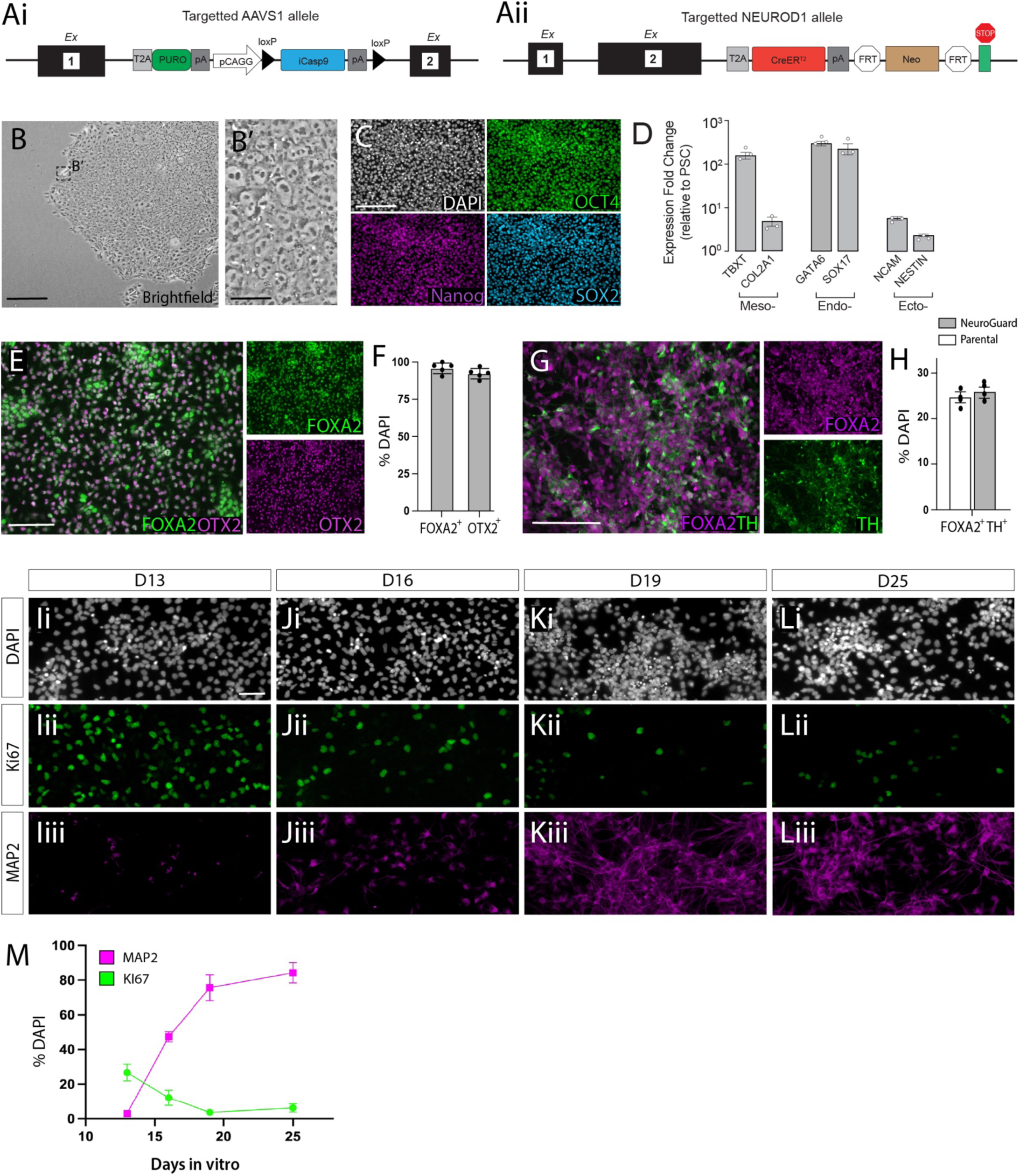
Design and validation of NeuroGuard pluripotent stem cell line, a suicide gene-based system designed to selectively protect neurons. (**Ai**) Schematic illustration of the targeting vector designed to insert the iCasp9 gene under the AAVS1 safe harbour gene promoter and, (**Aii**) a Cre recombinase cassette at the NeuroD1 locus. (**B**) Pluripotency of the new PSC line was validated by colony formation and morphology in addition to (**C**) high proportions of cells co-expressing the pluripotency markers OCT4, SOX2 and NANOG and, (**D**) trilineage differentiation potential with associated upregulation of meso-, endo- and ectoderm genes by qPCR. (**E**) Representative images and (**F**) quantification illustrating the capacity of the NeuroGuard PSC line to differentiate into vm progenitors, indicated by the high proportion of FOXA2^+^ and OTX2^+^ cells at 13 days *in vitro*. (**G**) Matured vm cultures immunolabelled for FOXA2 and TH (**H**) and cell counts, highlighting proportion of dopaminergic cells in culture after 25 days *in vitro*, comparable to the Parental IPSC line, RM3.5. (**I-L**) Representative images (**M**) and quantification of maturing vm cultures from Day 13 to 25, illustrating the increasing proportion of post-mitotic neurons, and converse reduction in Ki67^+^ proliferative cells. Data represents Mean ± SEM, n=3 independent differentiations/condition/cell line. Scale bars: (B,C) 500 μm; (B’,I-L) 50 μm; (E,G) 200 μm. Abbreviations: D, days *in vitro*; Ecto-, ectoderm; Endo-, endoderm; meso, mesoderm; TH, tyrosine hydroxylase.

The new hPSC ‘NeuroGuard’ line was validated to be karyotypically normal (data not shown) and pluripotency confirmed by colony formation and morphology in addition to the high proportion (>99%) of cells immunolabelled for OCT4, SOX2 and NANOG (***Figure 1B-C***). Multi-lineage potential of the NeuroGuard hPSC line was confirmed using the STEMdiff Trilineage Differentiation Kit (StemCell Technologies) and RT-qPCR for upregulated lineage-specific targeted genes (Mesoderm: TBXT/brachyury and COL2A1/collagen; Endoderm: GATA6 and SOX17; Ectoderm: NCAM and Nestin), ***Figure 1D***.

Prior to neural transplantation, the NeuroGuard hPSC line was assessed for (i) its ability to differentiate into ventral midbrain (vm) dopaminergic progenitors (suitable for transplantation), and (ii) the efficiency of the suicide system to ablate non-neuronal cells *in vitro*. NeuroGuard hPSCs competently differentiated into vm progenitors as verified by the high proportion of cells co-expressing the forebrain-midbrain protein, OTX2, and floorplate protein, FOXA2 (>95%) at Day 13 of differentiation, ***Figure 1E-F***. Ongoing maturation of the cultures to Day 25 confirmed their ability to generate tyrosine hydroxylase-expressing midbrain dopamine neurons, with comparable to efficiency to the RM3.5 Parental iPSC line***, Figure 1G-H,*** and that described by us and others ^12,13,26,33,34^.

Necessary to assess the efficiency of the NeuroGuard hPSC line to selectively protect neurons *in vitro* was an understanding of the maturation rate of the vm differentiation. As expected, a progressive decrease in Ki67^+^ proliferative progenitors was observed over time (Day 13: 26.69% ± 4.86; D25: 6.21% ± 2.37) and a reciprocal increase in MAP2^+^ post-mitotic neurons (Day 13: 4.22% ± 0.42; D16: 47.26% ± 2.93; D25: 84.24% ± 5.83), ***Figure 1I-M*.** Day 16 cultures represented an optimal stage in the differentiation when the cells were efficiently patterned to a vm fate and showed a high proportion of neurons (MAP2+>45%), yet retained a sufficient pool of proliferative progenitors to enable validation of off-target population ablation (i.e. non-neuronal cells) together with assessment of potential drug toxicity.

### Tamoxifen-induced Cre recombination selectively protects neurons in culture against CID-induce Caspase-mediated cell death

To assess the efficiency of the NeuroGuard suicide gene system, the hPSC line was differentiated into vm progenitors and treated at Day 13-16 with 4-hydroxytamoxifen (4-OHT, a potent, selective estrogen receptor modulator and active metabolite of tamoxifen) to drive Cre-recombinase, excising the iCasp9 suicide gene from all post-mitotic NEUROD1-expressing neurons in the culture. Subsequent administration of Chemical Inducer of Dimerization (CID, AP20187) at Day 15 was aimed at activating the iCasp9 suicide gene within non recombined cells (i.e. non-neuronal cells, not expressing NEUROD1), with culture composition finally assessed at Day 18, ***Figure 2A***.

**Figure 2.**
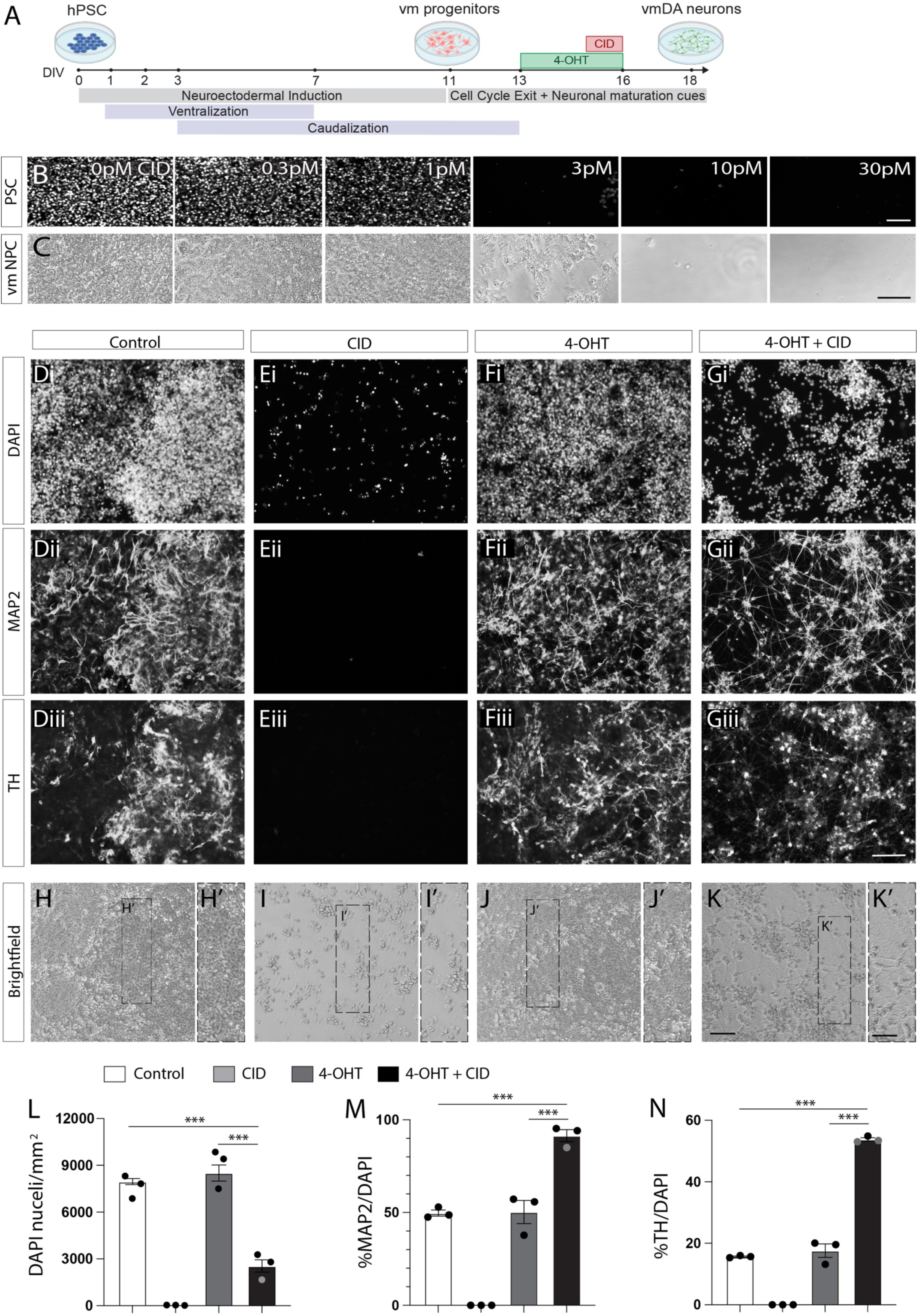
Activation of the NeuroGuard system selectively protects ventral midbrain dopamine neurons in culture. (**A**) Schematic illustrating vm DA differentiation and the timing of 4-OHT and CID treatment. (**B**) DAPI-labelled NeuroGuard PSCs and (**C**) brightfield images of vm progenitors treated with increasing doses of CID. Note, higher doses of CID are needed to ablate NPCs compared to PSCs. (**D**) Representative images of TH^+^ dopaminergic, MAP2^+^ neurons within D19 vm cultures showing (**E**) complete ablation following CID treatment and (**F**) no effect on 4-OHT cultures. (**G**) In contrast, 4-OHT+CID treatment showed less DAPI labelled cells, selectively protected only neurons in culture. (**H-K**) Brightfield images illustrating the maintained healthy morphology of neurons notably after 4-OHT+CID treatment (**K’**). (**L**) Quantification of total DAPI-labelled cells as well as (**M**) proportion of MAP2^+^ neurons and (**N**) TH^+^ dopamine neurons in untreated or 4-OHT and/or CID treated cultures. Note the significant increase in the proportion of neurons, and DA neurons, as a consequence of ablation of other cell types, namely progenitors in culture. Data represents Mean ± SEM, n=3 independent differentiations/condition. One-way ANOVA with Tukey’s multiple comparisons test. *p<0.05, **p<0.01, ***p<0.001. Scale bars: (B,C-K) 100 μm; (H’-K’) 50 μm. Abbreviations: CID, chemical inducer of dimerization; D, days *in vitro*; DA, dopamine; DAPI, 4’,6-diamidino-2-phenylindole; NPC, neural progenitor cells; PSC, pluripotent stem cells; TH, tyrosine hydroxylase; vm, ventral midbrain; 4-OHT, 4-hydroxytamoxifen.

Prior to assessment of the selective protection of neurons, a CID dose response confirmed successful expression of the iCasp9 gene within the NeuroGuard hiPSCs, with complete ablation of all undifferentiated cells achieved at 3pM (***Figure 2B***), whilst notably higher CID doses were required to similarly ablate vm neural progenitors/neurons (vm NPCs) within Day 16 vm cultures (***Figure 2C***), as previously observed in suicide gene based ablation of proliferative PSCs compared to NPCs ^26^.

Next, we compared untreated vm cultures, to those treated with CID and/or 4-OHT (***Figure 2D-K***). At Day 18, untreated cultures comprised 49.73% ± 1.65 MAP2^+^ neurons, inclusive of 15.67% ± 0.20 TH^+^ DA neurons, proportions that were unaffected by 4OHT treatment (50.27% ± 6.24 MAP2^+^; 17.62% ± 2.22 TH^+^), ***Figure 2M,N***. CID treatment ablated all cells in culture (***Figure 2E,L***). In contrast, 4-OHT treatment (from Day 13-16), followed by CID administration (at Day 15) resulted in a significant reduction in total cells in culture (65.44%, ***Figure 2L***) due to the ablation of non-neuronal cells, resulting in MAP2^+^ cells accounting for 91.48% ± 3.25 of total cells, and enrichment of TH+ neurons (53.77% ± 0.56), ***Figure 2G,M,N***.

### Establishing the maturity of human PSC-derived vm grafts

Studies of hPSC-derived neural grafts have demonstrated proliferation of vm progenitors after engraftment – inclusive of both the DA and non-DA fated populations ^17,20,26^. We therefore assessed the growth kinetics of the D19 vm progenitor grafts, derived from the new NeuroGuard cell line, with a focus on identifying the graft age at which the neuronal complement of the graft was maximal (as a proportion of total cells) and importantly the graft age at which the DA progenitors become post-mitotic neurons. Such knowledge will identify the optimal age to activate the selective ablation of non-neuronal cells within the graft – without affecting the functional unit of the graft (i.e. TH^+^ DA neurons) whilst ensuring the minimal proportion of cell death within the graft.

Volumetric assessment of grafts from 6 to 18 weeks post-transplantation revealed a steady increase in graft size, increasing >3-fold in size (6wks: 0.420 ± 0.082 mm^3^; 8wks: 0.474 ± 0.134 mm^3^; 10wks: 0.646 ± 0.186 mm^3^; 12wks: 1.036 ± 0.223 mm^3^; 18wks: 1.491 ± 0.371 mm^3^), ***Supplementary Figure 2Ai-Ei,-F***. Interestingly, between 12 and 18 weeks no significant increase in the proportion of NeuN cells was observed (6 weeks: 36.23% ± 2.52; 8 weeks: 47.75% ± 4.29; 10 weeks: 50.18% ± 3.54; 12 weeks: 60.90% ± 2.94; 18 weeks: 58.37% ± 2.27), with >40% of the graft comprised of non-neuronal cells, inclusive of proliferative neural and non-neural progenitors, other neural populations and off-target non-neural postmitotic cells, ***Supplementary Figure 2Aii-Eii,G***.

Critically, TH^+^ cell number was shown to peak 10 weeks post-transplantation (***Supplementary Figure 2Aiii-Eiii, H)***, suggestive that selective activation of the suicide gene in non-neuronal cells (inclusive of proliferative progenitors) at and beyond this time would not impact the dopamine contribution to the graft and thereby function.

### Treatment with 4OHT and CID selectively enriches the neuronal population in hPSC-derived vm progenitor grafts in the intact brain

With evidence of the full complement of TH^+^ DA neurons present within the vm progenitor-derived graft at 10 weeks, and in an effort to minimally destabilise the graft due to unnecessary/excessive apoptotic cell death of non-neuronal populations, we induced Cre-recombinase at the time corresponding to high proportion of NeuN^+^ cells within the graft (i.e. 10-12 weeks, ***Supp Fig 2G***), and prior to expoencial graft expansion (10 weeks, ***Supp Fig 2F***). Grafted animals were treated at week 10 with 4-OHT for 7 days, with CID administration on the final 2 days of 4-OHT treatment, ***Figure 3A***.

**Figure 3.**
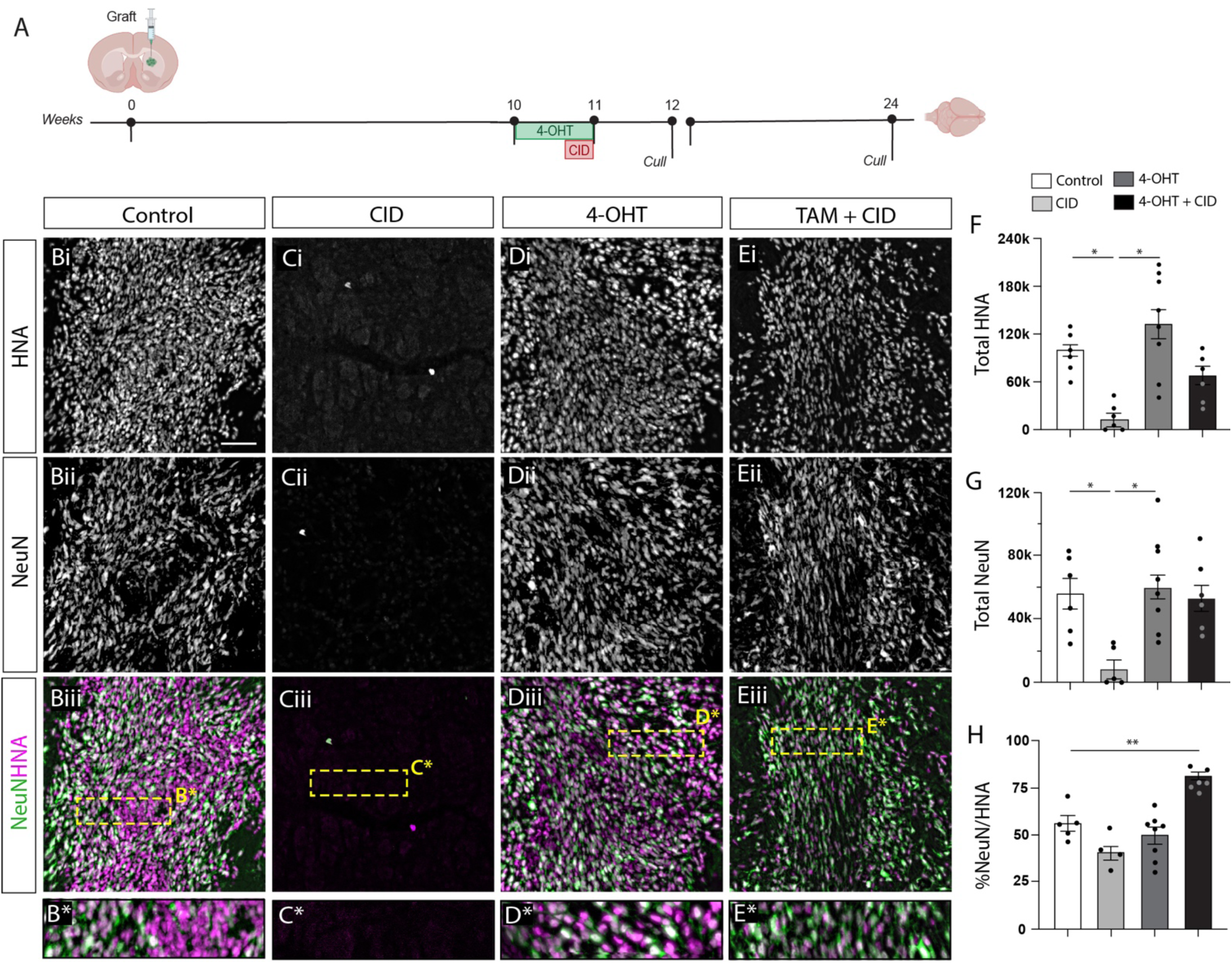
Activation of the NeuroGuard system selectively protects ventral midbrain dopamine neurons within neural transplants in the intact mouse brain. **(A)** Schematic illustration showing the timing of 4-OHT and CID treatment relatively to grafting in mice and the timing of postmortem analysis. (**B**) Representative images at 14 weeks depicting the proportion of NEUN^+^ neurons relative to total number of HNA^+^ cells within a graft in comparison to those animals treated with (**C**) CID, (**D**) 4-OHT or (**E**) 4-OHT + CID. Note the complete ablation of the graft after CID treatment. In comparison, untreated and 4-OHT treated mice showed grafts containing high neuronal pockets as well as numerous non-neuronal cells (NEUN^-^HNA^+^). 4-OHT+CID treated animals resulted in grafts that were almost entirely neuronal (NEUN^+^HNA^+^). (**F**) Quantification of total HNA^+^ cells, (**G**) NEUN^+^ cells and, (**H**) the relative proportion of neurons (%NEUN^+^HNA^+^/HNA^+^) within the grafts at 14 weeks after transplantation. Data represents Mean ± SEM, n=4-8 grafts/group. One-way ANOVA with Tukey’s multiple comparisons test. *p<0.05, **p<0.01, ***p<0.001. Scale bars: (B-E) 200 μm. Abbreviations: CID, chemical inducer of dimerization; HNA, human nuclear antigen; TH, tyrosine hydroxylase; wks, weeks4-OHT, 4-hydroxytamoxifen.

Assessment of grafts after 1 week (i.e. 12 weeks post-transplantation) revealed discrete transplants in all animals (***Supp Figure 3A-B***). In control, untreated mice, pockets of both graft-derived neurons (NeuN^+^HNA^+^: 55.82% ± 4.66) and non-neuronal populations (NeuN^-^HNA^+^) were observed, ***Figure 3B,H*.** CID treatment resulted in near complete ablation of the graft (***Figure 3C,F***), while 4-OHT Cre-recombination resulted in no difference compared to Control grafts (NeuN^+^HNA^+^: 50.10% ± 4.35, ***Figure 3D,F-H***). In animals treated with both 4-OHT and subsequently CID, ablation of non-neuronal cells in the graft resulted in a significant increase in the proportion of NeuN^+^HNA^+^ cells, accounting for the vast majority of cells within the graft (78.84% ± 2.41, ***Figure 3E,H***).

To ensure effective, stable ablation of non-neuronal cells in the graft, without evidence of emergence of quiescent stem cell-derived populations, an additional cohort of animals (treated with or without 4-OHT and CID at 10 weeks) was assessed at a protracted time frame (24 weeks post transplantation or 13 weeks after 4-OHT and CID treatment). As expected, in untreated animals graft size continued to increase (12 wks: 0.694 ± 0.075 mm^3^; 24 wks: 2.428 ± 1.147 mm^3^) and the proportion of NeuN^+^ cell remained relatively stable (***Supplementary Figure 3A,C,E,F).*** Reassured, in animals treated with 4-OHT and CID, no increase in graft size was observed (12 wks: 0.556 ± 0.103 mm^3^; 24 wks: 0.828 ± 0.145 mm^3^), with proportions of NeuN^+^ cells remaining comparable to 12 weeks (12 wks: 78.84% ± 2.41; 24 wks: 79.48% ± 3.23 %NeuN^+^/HNA^+^ cells) ***Supplementary Figure 2B,D,E,F, Figure 3H***.

### Treatment of grafted cells with 4-OHT and CID does not promote a significant immune response

With evidence of selective ablation of the non-neuronal cells within the graft following 4-OHT and CID administration, we next assessed the impact on local inflammation. Acutely after targeted apoptosis of the unwanted populations (12 weeks post-transplantation), animals showed no significant difference in the density of GFAP-immunoreactive astrocytes yet a 2.18-fold increase in microglial density of IBA1-immunoreactive pixels adjacent to the graft site, compared to untreated transplanted animals (Tx: 5.76 ± 2.53; Tx + 4-OHT + CID: 12.57 ± 1.65), ***Figure 4***. By 24 weeks post-transplantation (12 weeks after CID-induced targeted ablation), GFAP- and IBA1-immunoreactive density was not different between the treated and untreated animals, indicative of a transient microglial inflammatory response.

**Figure 4.**
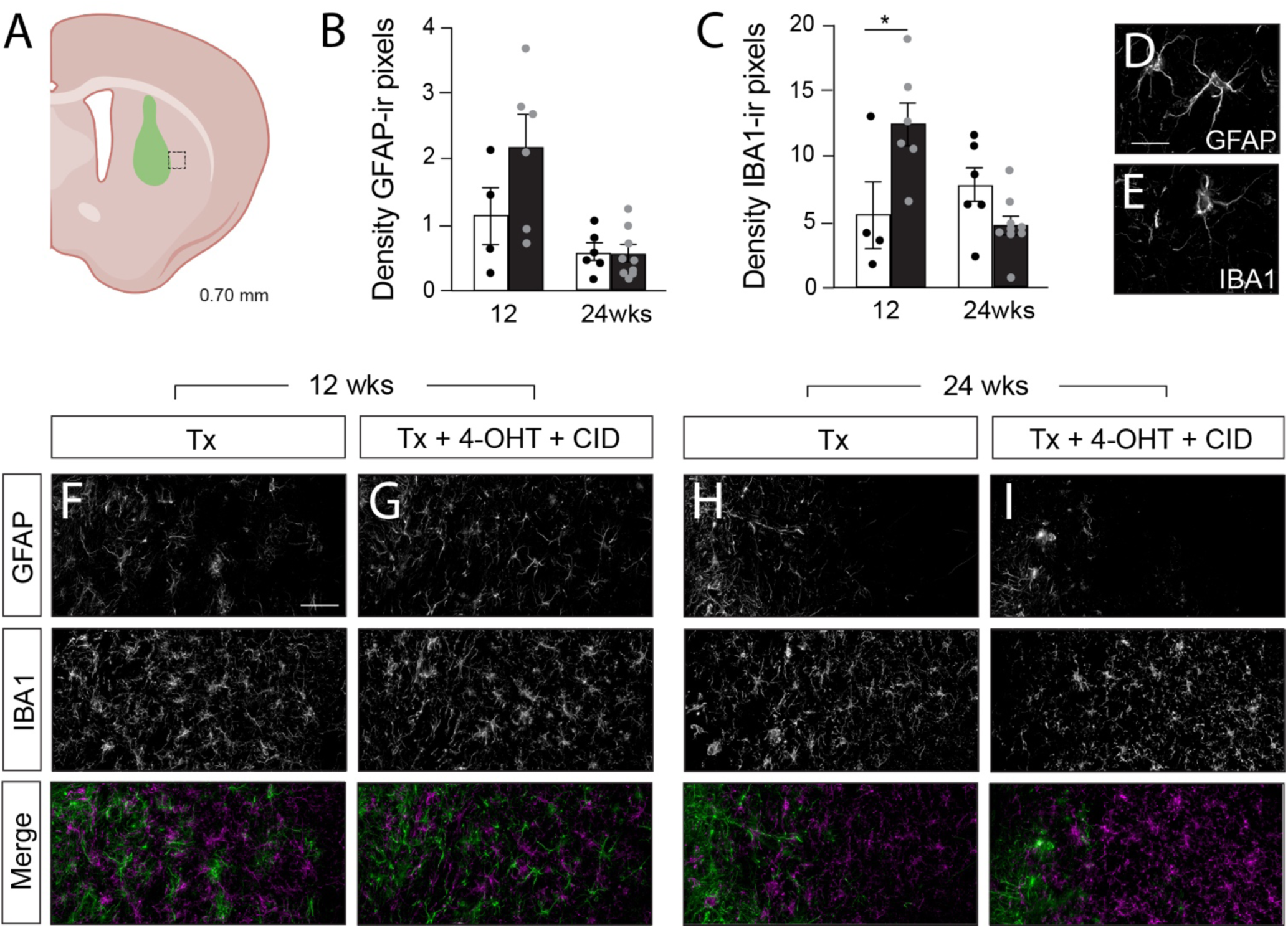
Activation of the NeuroGuard system in neural transplants transiently induces an microglial response. **(A)** Schematic illustrating area adjacent to the graft sampled for assessment of local inflammation. **(B)** Quantification of GFAP^+^ astroglial and, **(C)** IBA-1^+^ microglial responses at 14 and 24 weeks after transplantation - corresponding to 1 and 11 weeks after selective activation of the iCasp9 suicide gene. Data represented the density of GFAP or IBA-1 immunoreactive pixels as a proportion of total pixels. Data represents Mean ± SEM, n=4-6 grafts/group. Student t-test. *p<0.05. **(D)** Representative image of GFAP-labelled reactive astrocyte and, **(E)** IBA-1 stained microglia. **(F-G)** Example images depicting the density of astrocytes and microglia in untreated and 4-OHT+CID treated animals at 14 weeks and **(H-I)** 24 weeks after grafting. Scale bars: (**D,E**) 20 μm; (**F-I**) 100 μm. Abbreviations: CID, chemical inducer of dimerization; GFAP, Glial Fibrillary Acidic Protein; IBA-1, Ionized calcium-binding adapter molecule 1; ir, immunoreactive; Tx, transplantation; wks, weeks; 4-OHT, 4-hydroxytamoxifen.

While these findings highlight a transient inflammatory reaction in the host brain in response to targeted apoptotic ablation of non-neuronal populations following suicide gene activation, it is important to acknowledge the limitations of the model in which these studies are performed. Existing preclinical studies, relying on xenografting of human cells into rodent models (inclusive of athymic and immunosuppressive regimes) are unable to fully anticipate the immunological response to activation of the suicide gene system proposed here in the context of future human clinical allografts.

### Grafts treated with 4-OHT and CID integrate into a Parkinsonian rodent brain and successfully restore motor deficits

Next, we assessed the ability of NeuroGuard vm progenitor grafts to functionally integrate into a rat model of PD. Two weeks after 6-OHDA lesioning, amphetamine-induced rotational testing identified animals with motor deficits. Only animals exhibiting >300 rotations/hr were included in the study (totalling 22 of 30 rats) and stratified across 3 groups: (i) Lesion (n=7), (ii) Lesion + Transplant (Tx, n=7) and, (iii) Lesion + Transplant + 4-OHT + CID (Tx + 4OHT + CID, n=8). Rats received transplants of vm progenitors at 3 weeks post lesioning and a subgroup administered 4-OHT at 10 weeks (for 7 days) and CID for the final 2 days of 4-OHT treatment. Behavioural testing was repeated at 14, 18 and 22 weeks and finally animals culled at 24 weeks for histological analysis, ***Figure 5A***.

**Figure 5.**
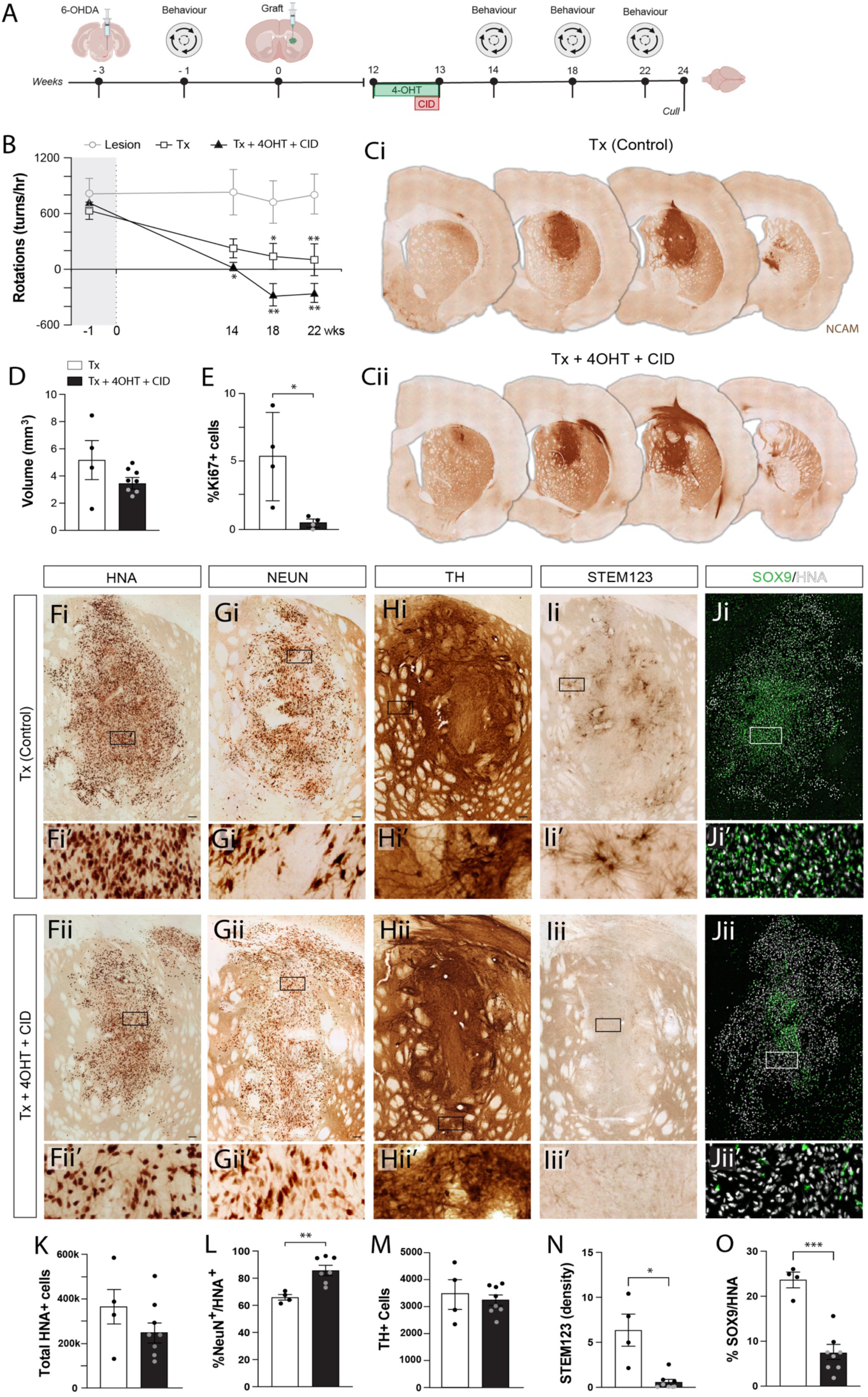
Activation of the NeuroGuard system in neural transplants in the Parkinsonian brain retains functional efficacy while eliminating off-target and proliferative cell populations. (**A**) Schematic illustration showing the timing of 6-OHDA lesioning, neural grafting, 4-OHT+CID treatment, behaviour and postmortem analysis in Parkinsonian rats. (**B**) Ablation of non-neuronal cells had no detrimental effect on graft function, with 4-OHT+CID treated rats showed full recovery of amphetamine-induced rotational asymmetry at 22 weeks. **(Ci)** Representative vm progenitor graft immunostained for NCAM and **(Cii)** in a PD rat that was treated with 4-OHT+CID. **(D)** Volumetric assessment of graft volume showed a visible, but not significant reduction in graft size at 24 weeks in 4-OHT+CID treated animals. **(E)** Critically, 4-OHT+CID treated animals showed a significant reduction in the presence of Ki67^+^ proliferative cells within neural grafts. **(F)** Representative images of NeuroGuard-derived neural graft from untreated and 4-OHT+CID treated PD rats illustrating the total number of graft-derived HNA^+^ cells, **(G)** NeuN^+^ neurons, **(H)** TH^+^ dopamine neurons, **(I)** human-specific STEM-123 labelled reactive astrocytes and **(J)** more broadly the proportion of total SOX9^+^ human astroglia. **(K)** Quantification of the total HNA^+^ cells in untreated and 4-OHT+CID treated rats and, **(L)** the relative proportion of NeuN^+^ neurons, noting the increase in neuronal purity. **(M)** Despite smaller grafts, containing few cells, TH^+^ cell counts remained unchanged. **(N)** Quantification of the number of STEM123 reactive astrocytes and proportion of SOX9^+^ cells revealed near complete ablation in 4-OHT+CID treated animals. Data represents Mean ± SEM, n=4-8 grafts/group. Student t-test. *p<0.05; **p<0.01; ***p<0.001. Scale bars: (C) 1mm; (F-J) 500 μm. Abbreviations: CID, chemical inducer of dimerization; HNA, human nuclear antigen; hr, hour; ir, immunoreactive; NCAM, neural cell adhesion molecule; TH, tyrosine hydroxylase; Tx, transplantation; wks, weeks; 4-OHT, 4-hydroxytamoxifen; 6-OHDA, 6-hydroxydopamine.

Importantly, the 6-OHDA-induced motor deficit was persistent and stable over the duration of the study (22 weeks) (Lesion, ***Figure 5B***, open grey symbols), with complete functional recovery in amphetamine-induced rotation observed by 18 weeks in animals receiving grafts without or with 4-OHT+CID treatment (black open and filled symbols, respectively, ***Figure 5B***).

Within the animal cohort two rats (Tx only group) were culled early due to respiratory disturbances and skin lesions (associated with long-term housing of immunocompromised rats). At 24 weeks post-transplantation, all animals showed surviving grafts, visible by human-specific NCAM expression within the host striatum (***Figure 5C***), with 1 animal excluded from the study due to poor graft placement (>50% the graft within the overlying cortex). Volumetric assessment of all remaining animals revealed that 4-OHT+CID treatment resulted in notably smaller grafts (Tx: 5.175 + 1.437 mm^3^; Tx + 4OHT + CID: 3.615 + 0.328 mm^3^, ***Figure 5D***), contained fewer cells, ***Figure 5D,F,K,*** and of visibly lower density (***Figure 5Fi’, Fii’***).

As previously observed the proportion of Ki67^+^ proliferative cells within the grafts at 6 months was low accounting for <5% of total HNA^+^ cells ^26,35^. Critically addressing safety, and a primary goal of the NeuroGuard system, this population was almost completely ablated upon 4-OHT + CID treatment, ***Figure 5E***.

Reflective of observations in mice, and validating efficiency of the NeuroGuard tool, quantitative assessment of neurons revealed a significant increase in their proportion in grafted rats treated with 4-OHT+CID (%NeuN/HNA, Tx: 65.89% + 2.02; Tx+4OHT+CID: 85.69% + 3.83), ***Figure 5G,L***. Because of neuronal protection, total TH^+^ dopamine neuron numbers within the grafts were not different between the grafted groups, complementing the functional recovery observed, ***Figure 5H,M***.

In light of the smaller grafts, we interrogated what populations were depleted from grafts treated with 4-OHT+CID. With previous studies showing the significant contribution of astrocytes within the vm transplants ^26^, we focused on this population. Human-specific reactive astrocytic marker STEM123, labelling for glial fibrillary associated protein, revealed a significant, 11-fold reduction in the glial population in 4-OHT+CID treated grafts, ***Figure 5J,N***. With STEM123 only labelling a subpopulation of astrocytes we additionally assessed the proportion SOX9^+^ cells within the grafts. Here SOX9 labelling could account for the majority of the non-neuronal cells in control, untreated grafts (%SOX9/HNA: 22.79 ± 1.77), yet was significantly reduced in those treated with 4-OHT+CID (%SOX9/HNA: 8.31 ± 1.97), ***Figure 5J,O***.

### Single nuclei transcriptional profiling confirms the elimination of non-neuronal populations within NeuroGuard transplants

To validate the ***in vivo*** profiling of hPSC-derived neural grafts with and without 4-OHT+CID treatment, we performed single nuclei transcriptional profiling on the same brains used for histological analysis using PathoSEQ^36^. Four grafts were pooled for both transplant conditions.

Quality control (***Supp Figure 4A)*** was performed across the pooled samples and highlighted comparable RNA quality across the nuclei of the two samples. Human nuclei were scored using gene sets representing human or rodent signature genes – eliminating Host (rodent), Human-Host doublets and any ‘Undetermined nuclei’, ***Supp Figure 4B***. Cluster analysis of the human nuclei from the two merged samples (treated and untreated) identified 9 populations, that could be further condensed to 7 populations based upon gene cluster analysis (***Supp Figure 4D, Figure 6Ai***). UMAP analysis of the 7 populations showed visually striking differences in the 2 groups – where neurons (including DA neurons) were highly enriched in Treated compared to Untreated grafts (82% and 28%, respectively), ***Figure 6Aii,iii***. This enrichment was at the expense of many other populations inclusive of radial glia/astrocytes, oligodendrocytes and neuroepithelial/neural stem cells that were markedly reduced in Treated grafts versus Untreated (15.2% and 50.5%, respectively), ***Figure 6Aii,iii***, supporting ***in vivo*** cell population counts (***Figure 5***).

**Figure 6.**
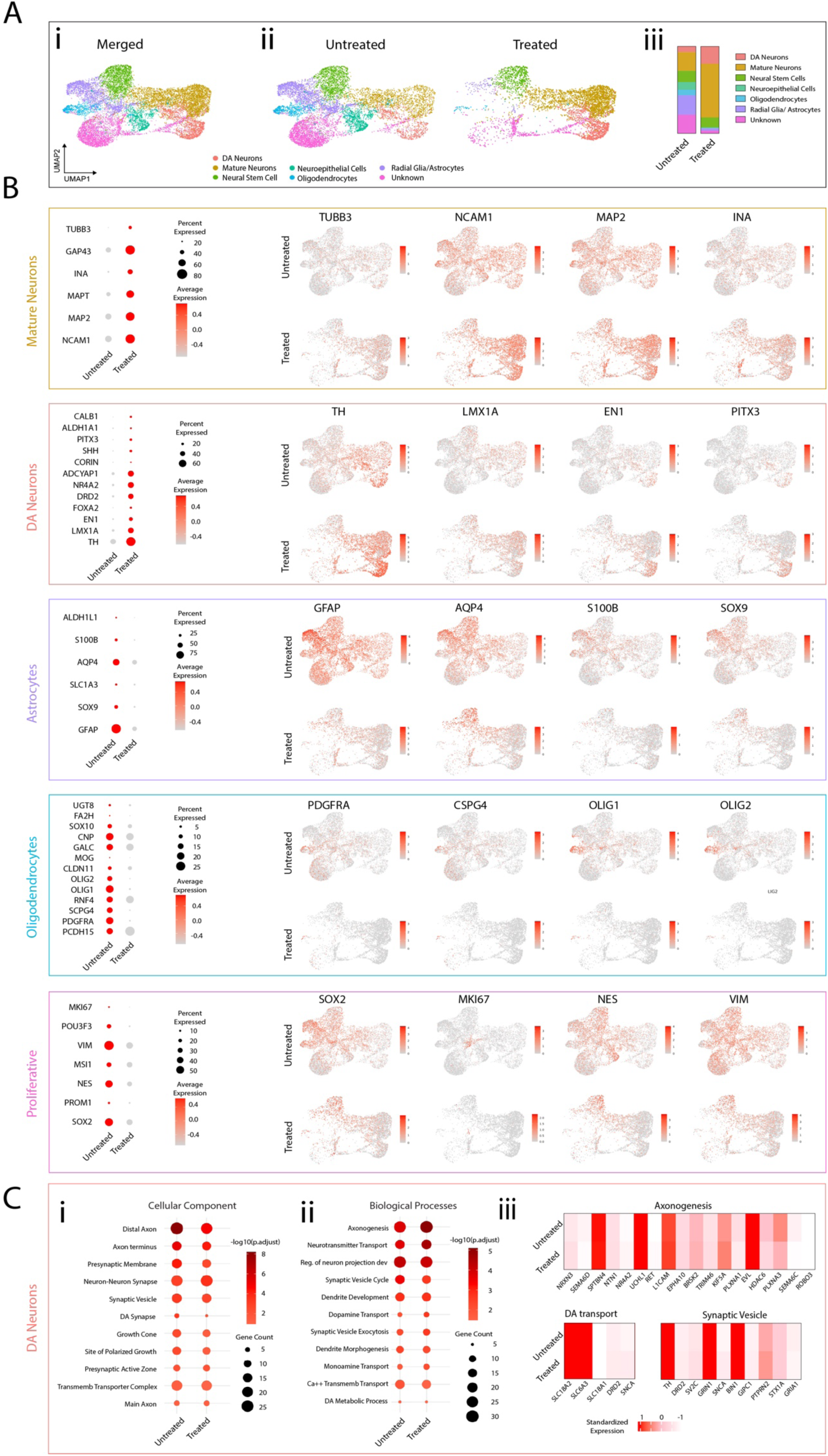
Single nuclei PathoSEQ reveals cell-type specific clustering with neuronal enrichment and their maintained maturity after 4OHT+CID selective ablation. **(Ai)** Human nuclei from the two samples were merged and based on initial Louvain clustering and scType identification annotated as DA neurons, mature neurons, neural stem cells, neuroepithelial cells, oligodendrocytes, radial glial/astrocytes and a ‘Mix’ of other indiscernible cell types. (**Aii**) UMAP and (**Aiii**) bar plot showing the proportion of different cell populations within mature grafts in untreated and 4-OHT+CID treated animals. **(B)** Bubble plot and UMAP showing the average expression, percent expression and distribution of key markers of the identified populations: mature neurons (TUBB3, NCAM1, MAP2), DA neurons (TH, LMX1A, EN, PITX3), astrocytes (GFAP, AQP4, S100B, SOX9), oligodendrocytes (PDGFRA, CSPG4, OLIG1, OLIG2) and proliferative cells (SOX2, Ki67, NES, VIM). **(C)** ‘GO ’terms for (**Ci**) cellular components and (**Cii**) Biological processes highlighting similarly upregulated pathways in grafted DA neurons in untreated and treated animals. (**Ciii**) Upregulated genes linked to axonogenesis, DA transport and synaptic vesicles showed a similar expression level in untreated and treated animals.

Importantly, differential gene analysis (and UMAP expression profiles), based on key markers (several included in histological analysis), verified the identity of the clusters as Mature Neurons (e.g. TUBB3, NCAM1, MAP2, INA), DA neurons (TH, LMX1A, EN1, PITX3), Astrocytes (GFAP, AQ4, S100β, SOX9),

Oligodendrocytes (PDGFRα, CSPG4, OLIG1/2) and Proliferative (SOX2, Ki67, NESTIN, VIMENTIN), ***Figure 6B***. Interestingly, in Untreated grafts 21.4% of the nuclei were classified as Unknown (compared to 3.1% in Treated grafts), representing a heterogeneous population that could not be ascribed a single fate (***Figure 6Aiii***, see pink proportion of bar).

Emerging studies have suggested that glial populations play an important role in the maturation of neural grafts^37–40^. Whilst our functional behaviour data suggested otherwise (***Figure 5B***), we nonetheless assessed the maturity of the dopamine neurons at a transcriptional level across the two treatment groups – Untreated Grafts (with high proportions of astrocytes) and Treated Grafts (with minimal astroglia present). Gene Ontology of Cellular Component and Biological Processes highlighted the remarkable similarity between the two groups, showing significant upregulation of gene clusters associated with ‘Neuron-Neuron Synapse’, ‘Synaptic Vesicle’, ‘Transmembrane Transporter Complexes’, ‘Axonogenesis’ and ‘Regulation of Neuron Projection Development’, (***Figure 6Ci,ii***). This was complemented by differential gene analysis highlighting conserved upregulation of axonogenesis genes such as L1CAM (previously identified in DA plasticity^41^), SPTBN4 (a critical voltage-gated sodium channels at the axon initial segment) and UCHL (necessary for DA neuron maintenance and survival) as well as DA transport and synaptic vesicle genes including SLC18A2/VMAT2 (essential for transporting dopamine into synaptic vesicles) , SLC6A3/DAT (essential for dopamine uptake from the synaptic cleft) and TH (tyrosine hydroxylase, essential for dopamine synthesis).

## DISCUSSION

The clinical translation of human pluripotent stem cell-derived cell therapies depends not only on advances in differentiation and manufacturing, but also on robust strategies to control graft composition, to ensure safety and maintain functional efficiency. Here we present NeuroGuard, a lineage-selective suicide gene platform that enables post-engraftment editing of transplanted cell populations. By selectively protecting post-mitotic neurons, whilst permitting inducible ablation of all other cell types, NeuroGuard addresses a central limitation of current approaches that fail to reconcile safety with therapeutic efficacy.

A defining feature of NeuroGuard is the ability to decouple safety from efficacy. To date, suicide gene strategies typically operate at one of two extremes to either eliminate proliferative cells while leaving grafts heterogeneous in composition ^24,26–29^, or entirely ablating the graft at the expense of its functional benefit ^7,30^. Recent advances in synthetic biology have highlighted the importance of multi-layered, programmable control systems, including logic-gated and inducible circuits, to regulate engineered cell behaviour *in vivo* ^8,31,32,42^. NeuroGuard extends this concept to regenerative medicine by introducing lineage-selective control over graft composition.

The NEUROD1-driven Cre recombination design of NeuroGuard enables the selective protection of the functional neuronal unit of the graft, rendering all other populations susceptible to apoptosis. This approach enables refinement of graft composition after transplantation without compromising function. This strategy aligns with emerging efforts to engineer next-generation cell therapies with built-in control mechanisms for safety, persistence and function ^6,8,42^.

In the context of a PD model and the transplantation of vm progenitors, here we show that NeuroGuard produces compact grafts with significantly increased neuronal proportion while preserving absolute numbers of dopaminergic neurons and maintaining full behavioural recovery. Smaller grafts with retained DA innervation capability reduce the displacement of the host tissue, while elimination of non-neuronal populations removes the ‘unknown’ regarding the potential impact of these populations on graft and host function. These findings demonstrate that therapeutic efficacy can be retained despite extensive removal of non-neuronal populations, while reducing graft size and heterogeneity - key parameters for clinical safety and scalability.

Another key finding of the study is that graft-derived astrocytes are dispensable for maintaining dopaminergic neuron survival and function at later stages of graft maturation. Previous studies have suggested that astrocytes enhance neuronal differentiation and functional integration ^37–39^, yet our data demonstrate that near-complete ablation of graft-derived astroglia does not impair neuronal survival, innervation or behavioural recovery. These findings suggest that host-derived glial populations may provide sufficient support for graft survival, integration and maturation.

Single-nuclei transcriptional profiling of the grafts further confirms the selective action of NeuroGuard, demonstrating enrichment of neuronal populations and depletion of astrocytic, oligodendroglial and progenitor lineages ^9,15,43,44^. Importantly, neuronal populations retained expression of genes associated with synaptic function, axonogenesis and dopamine neurotransmission, indicating preservation of functional identity and maturity. These findings demonstrate that post-engraftment editing of cell composition can be achieved without disrupting core transcriptional programs, a critical requirement for therapeutic application.

Beyond graft optimisation, safety remains a central barrier to the deployment of hPSC-derived therapies. While inducible suicide gene systems such as iCasp9 have demonstrated clinical utility^7^, they are not without limitations, including heterogeneous expression, incomplete killing and potential resistance ^27,45^. More broadly, the need for robust and fail-safe control systems has been highlighted across engineered cell therapy platforms. NeuroGuard mitigates some of these limitations by shifting from indiscriminate or proliferation-based targeting to lineage-selective control.

The selective ablation of cells within the brain raises concerns regarding host inflammatory responses. Here we observed a modest, transient increase in microglial density acutely after selective ablation of the non-neuronal cells, but no change in the density of reactive astrocytes. Importantly, this microglial response resolved over time and did not compromise DA neuron survival or functional recovery. While xenograft models cannot fully recapitulate immune responses in clinical allografts, these findings suggest that controlled, localised apoptosis within a graft can be accommodated by the host brain without deleterious consequences. One must however note the limitations of the xenograft model in immunodeficient rodents which cannot fully capture immune responses likely to occur following suicide gene activation in clinical allogeneic settings. Although the observed microglial response to targeted ablation was modest and transient, its magnitude may differ in patients.

The selection of NEUROD1 as the driver of neuronal protection is a key feature of the NeuroGuard platform. NEUROD1 is broadly expressed in post-mitotic neurons and plays essential roles in neuronal maturation and survival, including within vmDA populations ^46–49^. Targeting this developmental stage preserves a sufficiently large neuronal cohort to maintain graft stability while avoiding excessive ablation (as may occur through more restrictive promotors such as TH or GIRK2), that could destabilise graft architecture. Importantly, this strategy confers broad applicability beyond PD, positioning NeuroGuard as a versatile platform for other neuronal transplantation paradigms, including protecting medium spiny neurons in the treatment of Huntington’s disease, transplantation of interneurons for epilepsy or cortical neuron grafting for stroke. Added to this, the modular design of NeuroGuard raises the possibility of selectively protecting alternative cell types in other regenerative contexts through tailored promoter selection. In this regard, one may consider the prospects of selectively protecting only cardiomyocytes in transplants intended for cardiac repair or alternatively insulin-producing beta cells to control diabetes. In this context, NeuroGuard aligns with emerging efforts to develop programmable cell therapies with tuneable control over survival, identity and function, a defining goal of next-generation biotechnology platforms.

The prospects for this technology in the clinic in near future are real. Already the iCasp9 system has been clinically adopted to eliminate stem cell transplants where patients developed graft-versus host disease ^7^. Furthermore, both 4-OHT/tamoxifen and CID are already FDA approved and whilst earliest studies used FK506 as the FK binding partner, the newer chemical inducer of dimerization (CID) formulation, AP20187 employed in the present study, has been engineered to target the intrinsic apoptotic pathway and not interact with endogenous FKBP ^50^.

In summary, NeuroGuard establishes a first-in-class strategy for controlling the composition of transplanted cell populations *in vivo*. By enabling selective preservation of functional cells alongside targeted elimination of unwanted populations, this approach provides a powerful strategy to enhance the safety, predictability and scalability of cell-based therapies.

## MATERIALS & METHODS

### Generation of NEUROD1^CreERT^:AAVS1^loxP-iCasp9^ human PSC line

The human induced pluripotent stem cell (hiPSC) line, RM3.5, was maintained on Laminin-521 (0.5µg/cm2, Biolamina) as previously described^51^. To generate the new hPSC line, engineered to express iCaspase9 under the *AAVS1* promoter followed by subsequent editing at the *NEUROD1* locus to introduce a Cre recombinase cassette herein referred to as NEUROD1^CreERT^:AAVS1^loxP-iCasp9^.

Targeting of the *AAVS1* safe harbour in PSCs was achieved by the delivery of CRISPR-Cas expression and iCasp9 donor plasmids (Supp Figure 1B) by lipofection of pre-plated stem cells with these constructs.Selection with puromycin (PURO) (at 2μg/mL) from 6 days after transfection yielded sparse single-cell-derived clonal lines, at least 12 Puromycin-treated colonies were manually dissected and expanded for an additional 6 days. Following expansion, cells were isolated and genomic DNA (gDNA) was extracted using the ISOLATE II genomic isolation kit (Bioline). Resistant colonies were screened by RT-PCR for validation of the correct integration of the donor construct (Supp Figure 1C) into the *AAVS1* locus previously described ^52^ and the absence of extraneous targeting vector insertions ^40^

Successfully integrated clones displayed bands at 1312 bp following PCR of primers (1) & (2) (Supp Figure 1D). Confirmation of no extraneous targeting vector insertions was observed in clone #3 data not shown and was, therefore; further expanded to undergo a secondary round of targeting for insertion of conditionally activated Cre-recombinase at the *NEUROD1* locus. Clone #3 RM3.5: -loxP-iCasp9 PSCs were subjected to the identical process of genomic targeting by the delivery of CRISPR-Cas9 and CreERT2 custom-designed donor plasmids (Supp Figure 1E)) using lipofectamine.

Following transfection (120 hrs) of plasmids, positive selection of Neomycin-resistant clones was performed using G418 (10 μg/ml) and yielded considerable number of colonies. At least 20 colonies were manually passaged across 24- and 6-well plates for gDNA isolation and further expansion, respectively. Genomic DNA was isolated from these clones (as performed above) and RT-PCR was performed on 100 ng DNA/reaction with custom primers to amplify the genomic region flanked by donor homology arms confirming specific integration at the *NEUROD1* locus, Supp Figure 1F-G

Clones #8 and #17 demonstrated expected recombination of donor plasmid at the desired locus Supp Figure 1H. Based on intensity of cassette insertion and overall colony health, these clones were further expanded and underwent validation of iCasp9 induction by CID exposure and removal of this iCasp9 switch following TAM-mediated Cre-excision.

### Validation and ventral midbrain differentiation of the NEUROD1^CreERT2^:AAVS1^loxP-iCasp9^ hPSC line

Validation of pluripotency of the new *AAVS1*-iCasp9:*NEUROD1*-CreERT2 hPSC line was confirmed by immunocytochemistry (examining expression of OCT4, SOX2 and NANOG) as well as trilineage differentiation potential. Cultures of enzymatically-passaged PSCs were differentiated into the three main germ layers (ectoderm, endoderm and mesoderm) in monolayer culture using the STEMdiff Trilineage Differentiation Kit (StemCell Technologies). Differentiated cultures were harvested on day 5 (endoderm and mesoderm) or day 7 (ectoderm) for a total RNA extraction, using ISOLATE II RNA minikit (Bioline, UK). Expression of mRNA of distinct germ layer differentiation marker genes after differentiation was assessed by qPCR using lineage-specific targeted markers.

For generation of ventral midbrain progenitors/neurons, cells were differentiated according to the previously described protocol ^12,51^. In brief, cells were exposed to dual-SMAD inhibition using SB431542 (10μM, Tocris, days 0-5) and LDN193189 (200nM, Tocris, days 0-11). To ventralise the culture, Sonic hedgehog (SHH, 100ng/ml, R&D) and Purmorphamine (PM, 2μM, Tocris) were added from days 1-7. Wnt agonist CHIR99021 (2.5µM, Tocris) was added from days 2-13 to promote caudalisation. From day 11, cells were transitioned to a maturation media supplemented with BDNF (20ng/ml, R&D), GDNF (20ng/ml, R&D), TGFβ3 (1ng/ml, Peprotech), DAPT (10µM, Tocris), ascorbic acid (AA, 200μM, Sigma), and dibutyryl cAMP (dcAMP, 0.25mM, Sigma).

For *in vitro* validation of an active suicide gene, both undifferentiated hPSC and differentiation day 25 vm cultures were treated with increasing concentrations of Chemical Inducer of Dimerization, CID (AP20187, MedChem Express). To verify the selective excision of the suicide gene within neurons and protection thereof, day 13 vm differentiated cultures were treated with 1μg/ml 4-hydroxy tamoxifen (4-OHT, Sigma Aldrich) for 3 days, followed by 10pM CID (AP20187, MedChem Express) on day 16. Cells were then fixed 48h later for immunocytochemistry.

For *in vivo* transplantation studies, day 19 vm progenitors were dissociated in Accutase (StemCell Technologies) and resuspended into maturation media supplemented with ROCK inhibitor (Ri, 10μM, Sigma-Aldrich) at 100,000 cells/μl.

### In vivo procedures

#### Animal ethics statement

All animal procedures were performed in agreement with the Australian National Health and Medical Research Council’s published Code of Practice for the Use of Animals in Research, and approval granted by The Florey Institute of Neuroscience and Mental Health Animal Ethics committee. Animals (of either sex) were group housed on a 12:12h light/dark cycle with ad libitum access to food and water. All surgeries were performed under 2–5% isoflurane anaesthesia.

#### Surgeries

A total of 48 Rag1 knockout, immunodeficient mice and 30 adult athymic (CBH^rnu^) nude were used within the present study. Mice received stereotaxic intrastriatal cell grafts (0.5mm anterior, 2.5mm lateral to bregma, and 4.0mm below the dura surface) of Day 19 vm progenitors (100,000 cells in 1 μl).

To ablate the host midbrain dopamine neurons and pathways, athymic rats received unilateral injections (3.5 µl) of 6-OHDA (3.5 μg/μl free base dissolved in a solution of 0.2 mg/ml L-ascorbic acid in 0.9% w/v NaCl) into the medial forebrain bundle (MFB, 3.4mm posterior, and 1.3mm lateral to bregma, and 6.8mm below the dura surface dura surface) using a 10 ml Hamilton syringe fitted with a glass capillary. At 3weeks after 6-OHDA lesioning, following behavioural assessment, animals received unilateral cell grafts (1 µL, 100,000 cells/µl), into the denervated striatum (0.5 mm anterior, 2.5 mm lateral to bregma and 4.0 mm below the dura surface).

#### Tamoxifen and CID Treatment in vivo

For *in vivo* applications, 4-OHT was dissolved in ethanol to produce a 20mg/ml stock solution. On the day of injection, 1 part stock solution was mixed with 2 parts Chen oil (4 parts sunflower oil and 1 part castor oil). This mixture underwent vacuum centrifugation to evaporate the ethanol, resulting in a 10mg/ml solution. CID was dissolved in ethanol to produce a 50mg/ml stock solution. This was diluted to 0.4mg/ml (mice) or 3mg/ml (rats) in a solution of 4% ethanol, 10% PEG-400, and 2% Tween-20 in water on the day of injection. 10 weeks after transplantation, animals in the treatment group underwent daily intraperitoneal (IP) injections of 4-OHT (50mg/kg) for 7 days. CID (2.5mg/kg) was injected IP for the last 2 days of 4-OHT treatment. Mice were perfused 1 week following treatment. Rats were perfused at 24 weeks following behavioural assessment.

#### Behavioural analysis

Testing for rotational asymmetry was conducted 2 weeks after 6-OHDA administration to confirm motor deficits in unilateral lesioned rats. Net rotations over 60 minutes were analysed 10 minutes after intraperitoneal injection of D-amphetamine sulfate (5 mg/kg, Tocris Biosciences). Upon completion of initial testing, all animals displaying a functional deficit (>300 rotations in 60 min) were ranked in order of the percentage rotational asymmetry and evenly distributed across the three treatment groups: (i) Lesion only, (ii) Lesion + graft, (iii) Lesion + graft + 4-OHT and CID treatment. To assess functional integration of transplanted cells, behavioural testing was repeated at 14, 18 and 22-weeks post transplantation.

#### Tissue processing and immunohistochemistry

For confirmation of appropriate *in vitro* maturation and vm specification, cultures were fixed using paraformaldehyde (PFA, 4% w/v, 10 min) at varying time points during the differentiation (13, 16, 19, 25 DIV). Cells were blocked in a solution of 5% normal donkey serum, 0.3% v/v Triton-X100 (Amresco, USA), and 0.2% sodium azide in PBS-/- for 1 hour and then incubated with primary antibodies at 4°C overnight, followed by secondary antibody incubation for 2 hours.

For assessment of graft survival, composition and integration, animals were killed by an overdose of sodium pentobarbitone (100 mg/kg) at 6, 8, 10, 12, 18, or 24 weeks after transplantation and transcardially perfused with 4% PFA. Brains were cryosectioned (40 µM; 12 series) on a freezing microtome and immunohistochemistry was performed on free-floating brain sections. The tissue was incubated overnight with primary antibodies diluted in the blocking buffer described above. Detection of the primary–secondary antibody complexes was through peroxidase driven precipitation of di-amino-benzidine (DAB), or conjugation of a fluorophore. Secondary antibodies raised in donkey were applied for 2 h at room temperature for fluorescent detection. Chromogenic detection of antibody-DAB complex was carried out using biotin-conjugated secondaries and incubation with DAB (0.5 mg/ml, 3 min), which was precipitated by addition of 1% w/v H^2^O^2^. Fluorescently labelled sections were cover-slipped with fluorescent mounting media (Dako) and chromogenically labelled sections dehydrated in alcohol and xylene and cover-slipped with DePeX mounting media (BDH Chemicals). See ***Supplementary Table 1*** for a list of antibodies used.

#### Quantification

Brightfield images of chromogenic staining were captured using a Leica DM6000 microscope. Fluorescent images were captured on an EVOS M5000 (Invitrogen) or a Thunder (Leica). For i*n vitro* quantification of pluripotency, vmDA fate specification, and neural maturation, images captured at 20x magnification were used to quantify the total number of DAPI, OCT4, SOX2, NANOG, FOXA2, OTX2, TH, MAP2 and Ki67 immunolabelled cells. Quantification of viable (non-pyknotic) DAPI and immunolabelled cells was performed across three fields of view (FOV) per well, with three wells (technical replicates) per culture condition and repeated across three independent experiments for all conditions.

For *in vivo* assessment human-specific PSA-NCAM immunolabelling was used to delineate the graft boundaries and estimate graft volume according to Cavalieri’s principle. Images captured at 20x were used to quantify the total number of human nuclear antigen (HNA^+^), KI67^+^ proliferative cells, NEUN^+^ neurons and STEM123+ astrocytes within the graft. The total cell counts were made from 3 FOV per section across the rostro-caudal axis of the graft (from 1:12 series), spanning approximately 3-4 sections per graft. Quantification of tyrosine hydroxylase (TH^+^) DA neurons was performed on live images throughout the entire graft from a 1/12 series. Total numbers were estimated using graft volume. The density of host-derived GFAP^+^ astrocytes, and IBA1^+^ microglia were estimated as GFAP or IBA-1 immunoreactive pixels as a proportion of the total pixelsa djacent to the graft.

#### Single-nuclei RNA isolation, sequencing and bioinformatic analyses fixed samples (PathoSeq)

The protocol for single nuclei isolation of fixed tissue was adapted from Wang *et al.,* 2024 ^36^. In brief, the human graft was visibly identifiable and dissected using a punch biopsy needle (2mm, KAI-700, 21056-KA) from serial brain slices of previously perfused-fixed mouse brain (see above). Grafts from x animals per condition were pooled together in a 1.5ml Eppendorf tube, with 100μl of Digestion buffer (PBS-/- with 1mg/ml Liberase TH (Roche Diagnostics), 0.2U/μl RNase Inhibitor, 0.5mM CaCl^2^) and mechanically homogenized using a Biomasher II (Biostrategies), with 30 strokes. Tissue was then digested in a total of 1ml of digestion buffer (37°C, 45 min, 800rpm) on a Thermomixer, 400μl of EZ buffer added to the sample (EZ Lysis Buffer with 1% BSA, 0.2U/μl of RNase inhibitor), sample mixed by inversion 5x and centrifuged for 5 min at 900g at 4°C. Supernatant was removed and pellet resuspended in 600μl of EZ buffer, mixed 5x by pipetting, incubated 5 min on ice, passed through a 25G needle (5-10x) and washed with 600μl of wash buffer (PBS-/- with 1%BSA and 2.5mM MgCl^2^, 0.2U/μl RNase inhibitor). The tissue was then pelleted by centrifugation (4°C, 900g, 5 min), resuspended in 1000μl of wash buffer gently using a p1000, pipetting 30x, filtered (70μm and 40μm PluriStrainer Mini filter). Single nuclei libraries were constructed using the GEM-X Flex Gene Expression kit for multiplex samples (10x Genomics), following the 10x user guide (CG000787_GEM-X_Flex_MultiplexedSamples_UserGuide_Rev_B). Nuclei were quantified using the LUNA-FL dual fluorescent cell counter and equal numbers of nuclei were combined for each sample. Libraries were sequenced on NextSeq2000 (Illumina) P4 flow cells, 100 cycles. Reads were demultiplexed and mapped to the reference genome (GRCh38-2024-A, build 2024-A, 10x Genomics) using Cell Ranger v9.0.1 (10x Genomics). Single-nuclei data were processed using Seurat ^53^ environment in R programming language. For each sample, high quality single-nuclei were retained that had < 10% mitochondrial counts, between 500 and 10000 features and more than 1000 RNA counts (**Supplementary Figure 4A**). Samples were then normalized, scaled and PCA, UMAP and clustering performed using Seurat inbuilt functions NormalizeData, ScaleData, RunPCA, RunUMAP ^54^. The dimension used for “FindNeighbors” was determined using “ElbowPlot.” Graft and host signature (**Supplementary Figure 4B**) was derived from the top and bottom of the first principal component of a dataset with known human and rat brain snRNA-seq (see list of genes on github https://github.com/parishgroup/iCasp9_snRNAseq/blob/main/data/PDgraft_ku80sorted_unsorted_P rincipleComponents.numbers ) and the subset of human nuclei denoted as “human”. Human nuclei from each sample were then merged ^55^, integrated normalized and clustered as described above. Cluster analysis of the human nuclei from the two merged samples identified 9 distinct populations, represented across the samples with different numbers of nuclei for each sample (**Supplementary Figure 4C, D**). To evaluate the cell-types of the human nuclei we used a reference-based single-cell RNA-seq annotation tool (scType)^56^ (**Supplementary Figure 4D**). Differential expression analysis was performed using Seurat function FindMarkers, followed by gene ontology enrichment (for the upregulated genes, with avg_log2FC > 0.25, pct.1 > 0.5, p_val_adj < 0.05) using the clusterProfiler package^57^ part of Bioconductor Gentleman et al., 2004 ^58^), with the org.Hs.eg.db annotation database (Carlson Bioconductor annotation package, DOI: 10.18129/B9.bioc.org.Hs.eg.db). GO terms from "GO_Molecular_Function" and "GO_Cellular_Component" databases were used^59^. Genes in the identified GO were visualized in a bar plot (**Supplementary figure 4E**) using the ComplexHeatmap package ^60^ (**Figure 6C**). The code used is available on github https://github.com/parishgroup/iCasp9_snRNAseq.

## Statistical analysis

All behavioural testing and histological assessments were performed with the researcher blinded to the experimental group. All data is presented as mean ± SEM. Statistical tests employed (one-way ANOVA, two-way ANOVA, Student’s t tests), and number of animals/groups for each assessment are stated in the figure legends. All statistical analyses were performed using GraphPad Prism and alpha levels of p<0.05 were considered significant (*p<0.05, **p<0.01, ***p<0.001).

## Data availability

The authors declare that the data supporting the findings of this study are presented within the paper and its supplementary information files. Source data are provided with this paper.

## Acknowledgements

The authors thank Emily Hart and Brianna Xuereb for expert technical assistance. JJ was supported by an was supported by an Australian government Research Training Program (RTP) stipend. CLP was supported by a National Health and Medical Research Council Australia (NHMRC) Leader Fellowship (level 2, GNT2026395). NM was supported by a postdoctoral fellowship provided by the EJ Flack Foundation, Australia. This work was supported by funding from the CASS Foundation, Australia. The Florey Institute of Neuroscience and Mental Health acknowledges the strong support from the Victorian Government and the funding from the Operational Infrastructure Support Grant. The authors thank Mr David Yoannidis and the Molecular Genomics Core at the Peter MacCallum Cancer Center for their technical support on the library preparation and sequencing.

## Author contributions

Conceptualisation: CLP. Experimentation: JJ, CP, CJPH, BX, NM, GF, ATQ, DO, CLP. Data analysis: JJ, CP, CPJH, NM, CLP. Contributed reagents: CLP. Manuscript preparation: JJ, CLP. Manuscript editing: JJ, CP, CJPH, GF, NM, DO, ATQ, CLP.

---

**Supplementary Figure 1.**
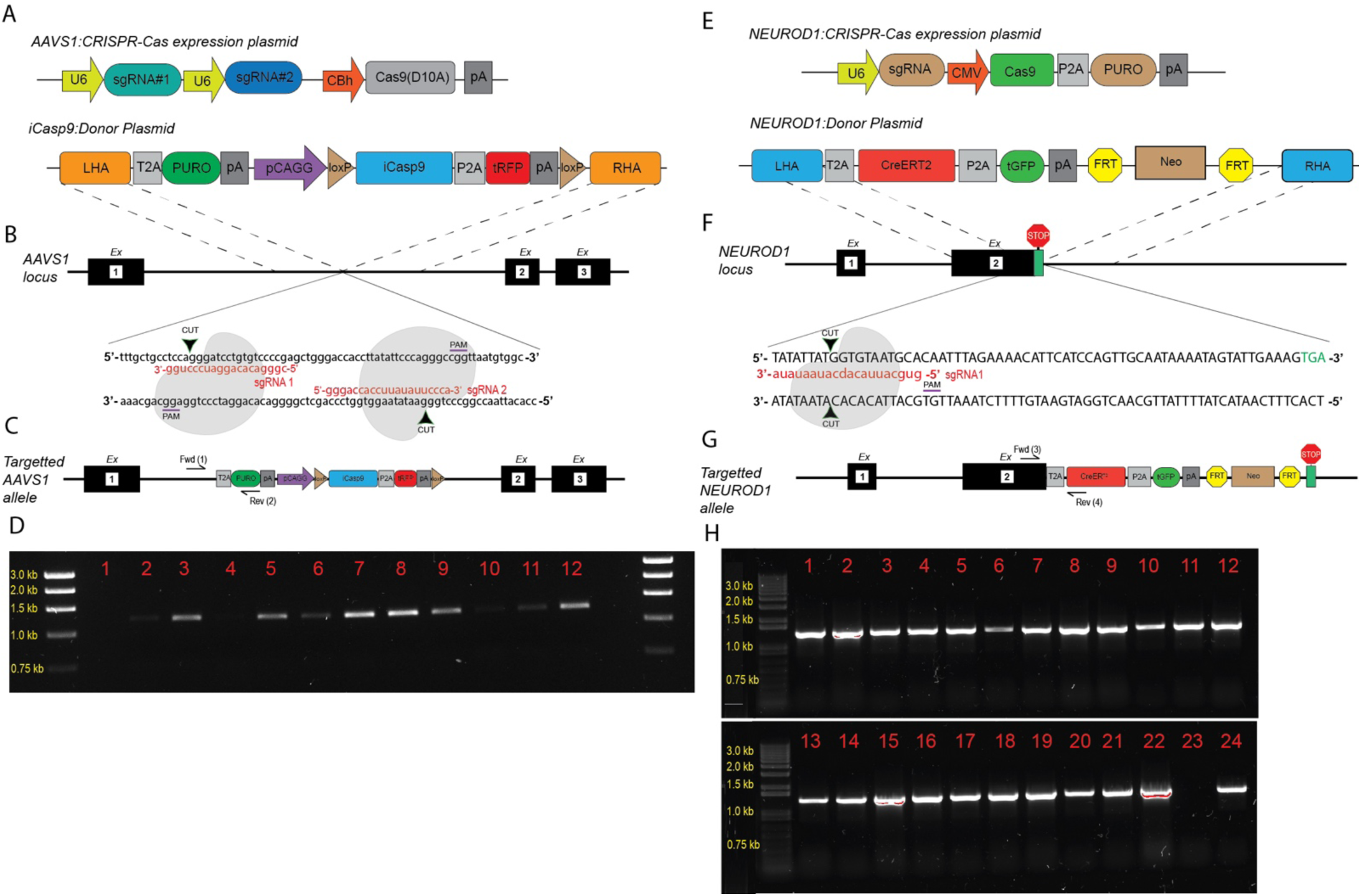
Sequential site-specific targeting of iCasp9 and CreERT2 ORFs to AAVS1 and NEUROD1 loci in PSCs. (A & E) Schematic representation of donor and CRISPR-Cas expression plasmids designed for site-specific integration at the AAVS1 and NEUROD1 loci. (B & F) Genomic detail of the AAVS1 and NEUROD1 loci indicating guide RNA target sites and regions of homology for recombination. (C & G) Predicted integration of donor plasmids and PCR strategy following correct recombination at AAVS1 (C) and NEUROD1 (G) loci. (D & H) PCR-validation of targeted integration using locus-specific primers. Representative agarose gels show expected band sizes for correctly targeted clones (red numbers) for AAVS1 and NEUROD1 regions, confirming site-specific integration.

**Supplementary Figure 2.**
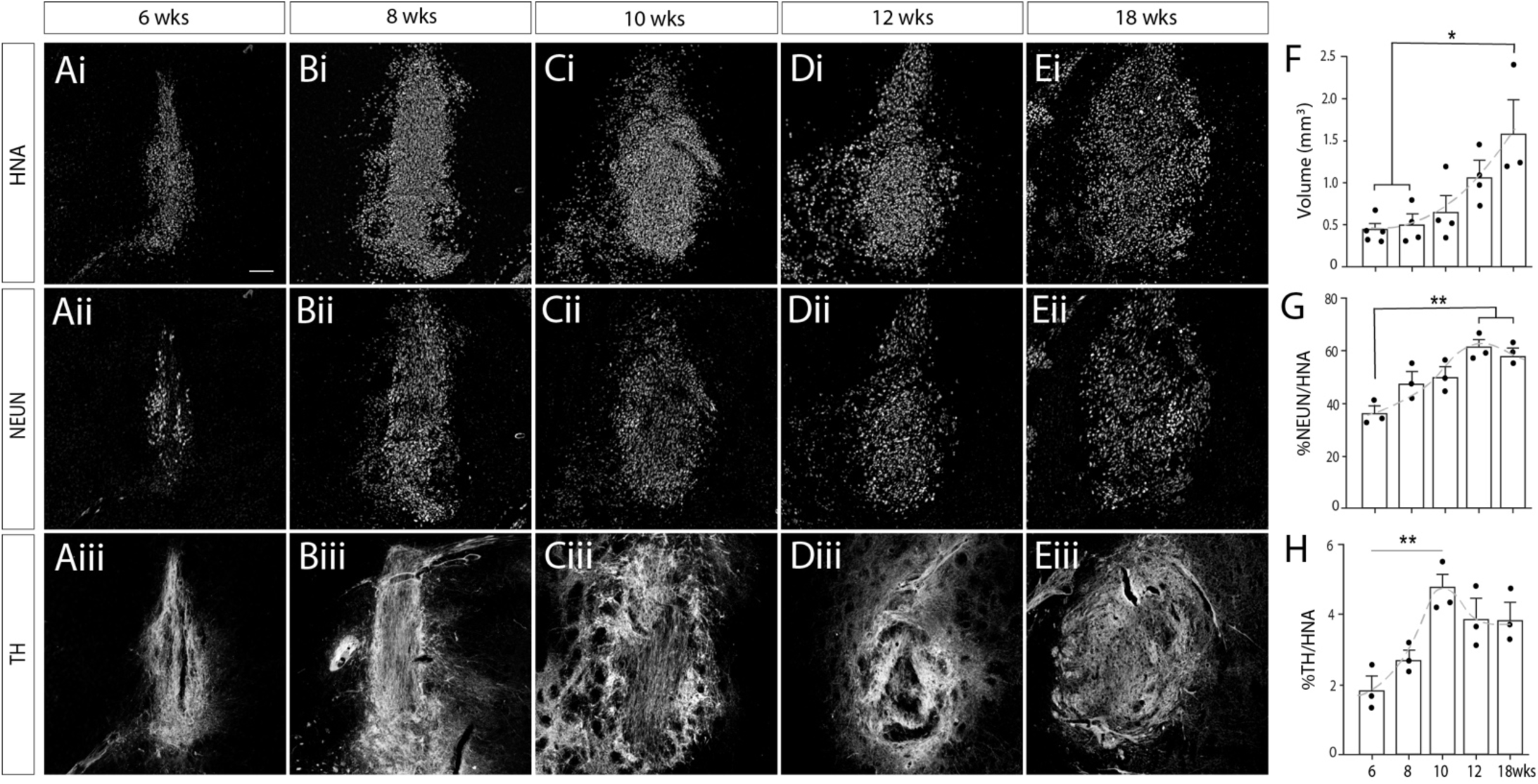
Understanding the kinetics of human PSC-derived vm progenitor graft maturation. (**A-E**) Representative images depicting the maturation of human PSC-derived vm progenitor grafts from 6-18weeks after implantation, indicating graft size (HNA) as well as the proportion of total neurons (NEUN) and more specifically dopaminergic (TH+) neurons. (**F**) Quantification of graft volume and, (**G**) the proportion of NEUN+ human neurons and, (**H**) TH+ dopamine neurons in the graft. Note the steady increase in graft volume with time yet plateau in TH+ cells as approximately 10-12 weeks after transplantation. Data represents mean + SEM, n=4-5 grafts/time point. Scale bars: (**A-E**) 200 μm. Abbreviations: HNA, human nuclear antigen; TH, tyrosine hydroxylase; wks, weeks.

**Supplementary Figure 3.**
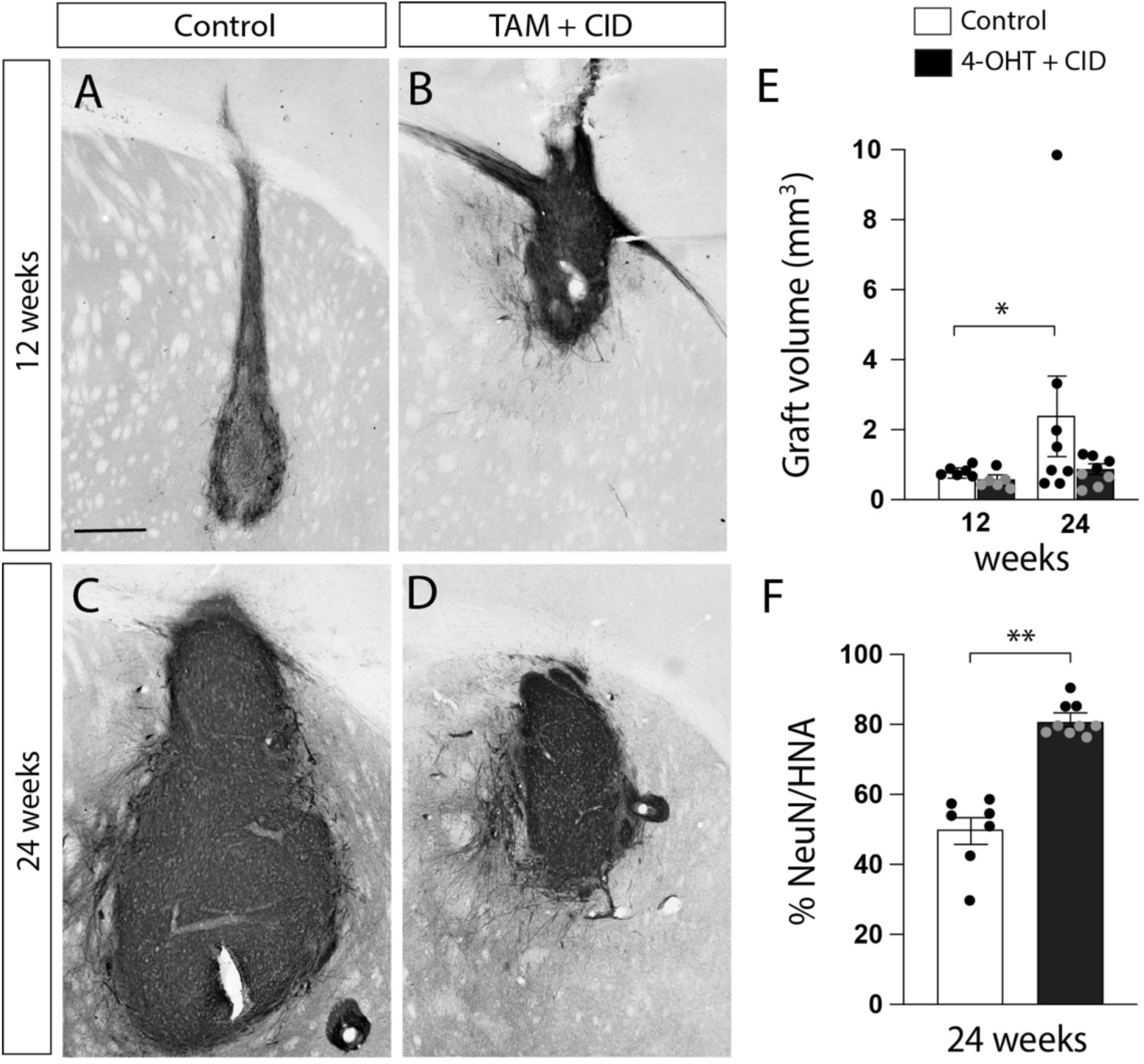
Activation of the NeuroGuard suicide system results in stable and effective ablation of non-neuronal populations. (**A-D**) Representative images showing human-specific NCAM labeling of the graft, illustrating the increase in graft size over time in Control animals, yet maintained discrete grafts in animals receiving 4OHT+CID treatment. (**E**) Quantification of graft volume and, (**F**) the proportion of NEUN^+^ human neurons in the graft at 12 and 24 weeks, highlighting that 4-OHT+CID treated grafts remained almost entirely neuronal (NEUN^+^HNA^+^). Data represents Mean ± SEM, n=6-8 grafts/time point. Student t-test. *p<0.05, **p<0.01, Scale bars: (**A-D**) 200 μm. Abbreviations: 4-OHT, 4-hydroxytamoxifen; CID, chemical inducer of dimerization.

**Supplementary Figure 4.**
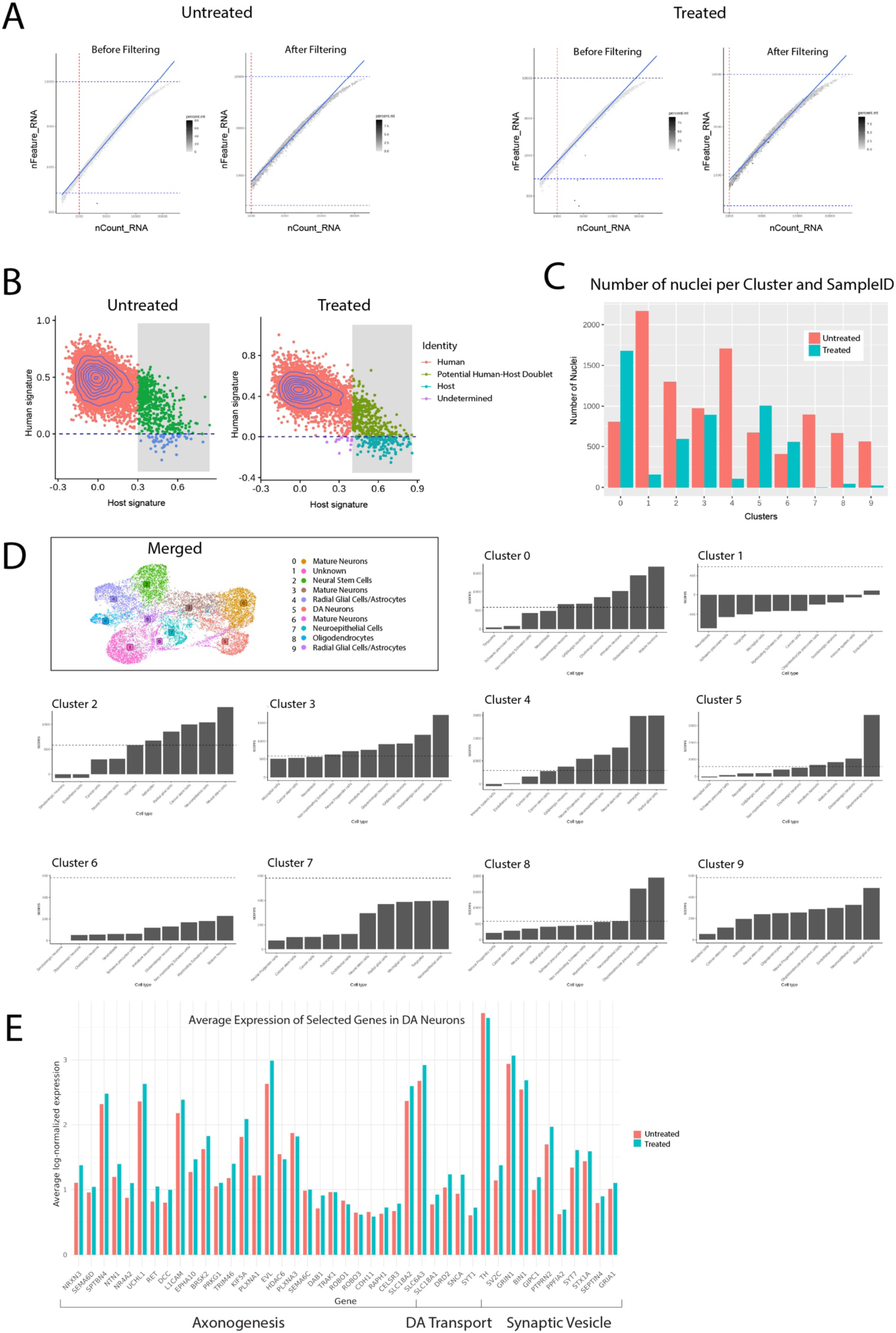
Single nuclei sequencing quality control and annotation. **(A)** QC performed for untreated and treated samples; only high quality single-nuclei were retained that had < 10% mitochondrial counts (grey color scale), between 500 and 10000 features (y axis nFeature_RNA) and more than 1000 RNA counts (x axis, nCount_RNA). **(B)** Human and rat signature across the two different samples. **(C, D)** Cluster analysis of the human nuclei from the two merged samples identified 9 distinct clusters, represented across the samples with similar number of nuclei for each sample. The identity of each cluster was identified using scType score. **(E)** Average express of selected genes in DA neurons showed similar expression in untreated and treated samples.

**Supplementary Table 1.**

| Target | Host | Supplier | Cat# | Dilution |
| --- | --- | --- | --- | --- |
| DAPI | - | Sigma Aldrich | D9542 | 1:5000 |
| FOXA2 | Mouse | Abcam | Ab60721 | 1:4000 |
| FOXA2 | Goat | Santa Cruz | sc-6554 | 1:200 |
| GFAP | Chicken | Invitrogen | PA11004 | 1:1000 |
| GFP | Chicken | Abcam | Ab13970 | 1:1000 |
| GFP | Rabbit | Abcam | Ab290 | 1:200 |
| Human nuclear antigen (HNA) | Mouse | Millipore | MAB1281 | 1:300 |
| IBA1 | Rabbit | Wako | 019-19741 | 1:1000 |
| KI67 | Rat | Invitrogen | 14-5698-80 | 1:300 |
| Nanog | Rabbit | Proteintech | 14295-1-AP | 1:200 |
| MAP2 | Chicken | Invitrogen | PA1-1005 | 1:2000 |
| NCAM |  |  |  |  |
| NeuN | Rabbit | Abcam | Ab236869 | 1:1000 |
| NeuN | Rabbit | Abcam | Ab104225 | 1:200 |
| OCT4 | Mouse | Santa Cruz Biotech. | SC5279 | 1:100 |
| OTX2 | Rabbit | Proteintech | 13497-1-AP | 1:200 |
| OTX2 | Goat | R&D Systems | AF1979 | 1:1000 |
| PSA-NCAM (Eric1) | Mouse | SantaCruz Biotechnology | sc-106 | 1:500 |
| PSA-NCAM (Eric1) | Rabbit | Abcam | Ab75813 | 1:1000 |
| SOX2 | Goat | R&D Systems | AF2018488 | 1:300 |
| SOX9 | Rabbit | Abcam | Ab185966 | 1:500 |
| STEM123 | Mouse | Takara | Y40420 | 1:500 |
| TH | Mouse | ThermoFisher | TA506549 | 1:500 |
| TH | Rabbit | Pel-freeze | P40101-0 | 1:1000 |

**Supplementary Table 2.**
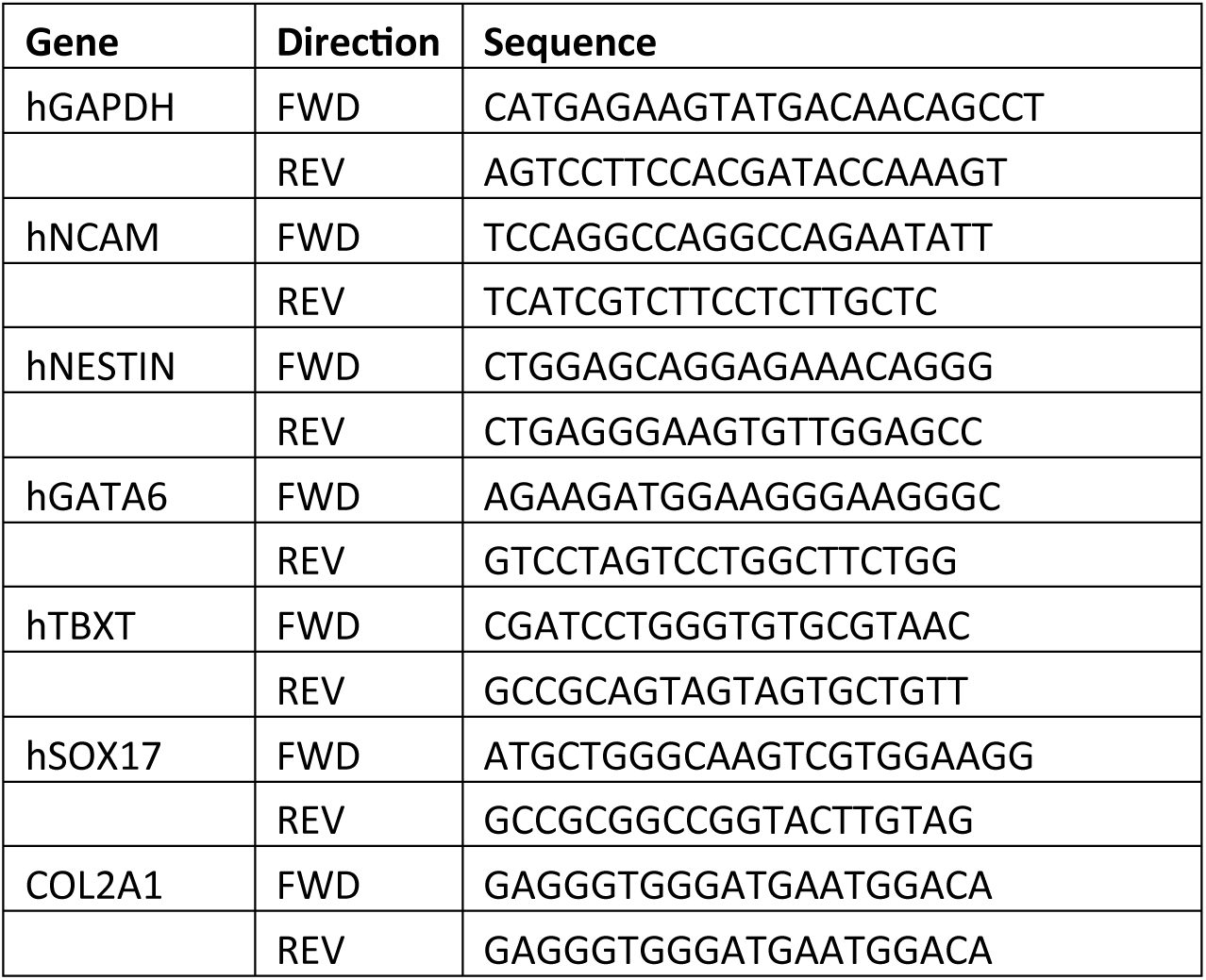

## REFERENCES

1 Kirkeby, A., Main, H. & Carpenter, M. Pluripotent stem-cell-derived therapies in clinical trial: A 2025 update. Cell Stem Cell 32, 329–331 (2025). 10.1016/j.stem.2025.01.003

2 Schweitzer, J. S. et al. Personalized iPSC-Derived Dopamine Progenitor Cells for Parkinson’s Disease. N Engl J Med 382, 1926–1932 (2020). 10.1056/NEJMoa1915872

3 Tabar, V. et al. Phase I trial of hES cell-derived dopaminergic neurons for Parkinson’s disease. Nature 641, 978–983 (2025). 10.1038/s41586-025-08845-y

4 Sawamoto, N. et al. Phase I/II trial of iPS-cell-derived dopaminergic cells for Parkinson’s disease. Nature 641, 971–977 (2025). 10.1038/s41586-025-08700-0

5 Barker, R. A., Parmar, M., Studer, L. & Takahashi, J. Human Trials of Stem Cell-Derived Dopamine Neurons for Parkinson’s Disease: Dawn of a New Era. Cell Stem Cell 21, 569–573 (2017). 10.1016/j.stem.2017.09.014

6 Kimbrel, E. A. & Lanza, R. Current status of pluripotent stem cells: moving the first therapies to the clinic. Nat Rev Drug Discov 14, 681–692 (2015). 10.1038/nrd4738

7 Di Stasi, A. et al. Inducible apoptosis as a safety switch for adoptive cell therapy. N Engl J Med 365, 1673–1683 (2011). 10.1056/NEJMoa1106152

8 Roybal, K. T. et al. Engineering T Cells with Customized Therapeutic Response Programs Using Synthetic Notch Receptors. Cell 167, 419–432 e416 (2016). 10.1016/j.cell.2016.09.011

9 Zhang, X. et al. 3D-generation of high-purity midbrain dopaminergic progenitors and lineage-guided refinement of grafts supports Parkinson’s disease cell therapy. Cell Stem Cell 32, 1758–1772 e1758 (2025). 10.1016/j.stem.2025.10.001

10 Kirkeby, A. et al. Generation of regionally specified neural progenitors and functional neurons from human embryonic stem cells under defined conditions. Cell Rep 1, 703–714 (2012).

11 Kriks, S. et al. Dopamine neurons derived from human ES cells efficiently engraft in animal models of Parkinson’s disease. Nature 480, 547–551 (2011).

12 Niclis, J. C. et al. Efficiently Specified Ventral Midbrain Dopamine Neurons from Human Pluripotent Stem Cells Under Xeno-Free Conditions Restore Motor Deficits in Parkinsonian Rodents. Stem Cells Transl Med 6, 937–948 (2017).

13 Kim, T. W. et al. Biphasic Activation of WNT Signaling Facilitates the Derivation of Midbrain Dopamine Neurons from hESCs for Translational Use. Cell Stem Cell 28, 343–355 e345 (2021). 10.1016/j.stem.2021.01.005

14 Kikuchi, T. et al. Human iPS cell-derived dopaminergic neurons function in a primate Parkinson’s disease model. Nature 548, 592–596 (2017). 10.1038/nature23664

15 Tiklova, K. et al. Single-cell RNA sequencing reveals midbrain dopamine neuron diversity emerging during mouse brain development. Nat Commun 10, 581 (2019). 10.1038/s41467-019-08453-1

16 Rajova, J. et al. Deconvolution of spatial sequencing provides accurate characterization of hESC-derived DA transplants in vivo. Mol Ther Methods Clin Dev 29, 381–394 (2023). 10.1016/j.omtm.2023.04.008

17 Niclis, J. C. et al. A PITX3-EGFP Reporter Line Reveals Connectivity of Dopamine and Non-dopamine Neuronal Subtypes in Grafts Generated from Human Embryonic Stem Cells. Stem Cell Reports 9, 868–882 (2017).

18 de Luzy, I. R. et al. Isolation of LMX1a Ventral Midbrain Progenitors Improves the Safety and Predictability of Human Pluripotent Stem Cell-Derived Neural Transplants in Parkinsonian Disease. J Neurosci 39, 9521–9531 (2019). 10.1523/JNEUROSCI.1160-19.2019

19 Doi, D. et al. Isolation of human induced pluripotent stem cell-derived dopaminergic progenitors by cell sorting for successful transplantation. Stem cell reports 2, 337–350 (2014). 10.1016/j.stemcr.2014.01.013

20 Samata, B. et al. Purification of functional human ES and iPSC-derived midbrain dopaminergic progenitors using LRTM1. Nat Commun 7, 13097 (2016).

21 Aguila, J. C. et al. Selection Based on FOXA2 Expression Is Not Sufficient to Enrich for Dopamine Neurons From Human Pluripotent Stem Cells. Stem Cells Transl Med 3, 1032–1042 (2014). 10.5966/sctm.2014-0011

22 Lehnen, D. et al. IAP-Based Cell Sorting Results in Homogeneous Transplantable Dopaminergic Precursor Cells Derived from Human Pluripotent Stem Cells. Stem Cell Reports 9, 1207–1220 (2017). 10.1016/j.stemcr.2017.08.016

23 Li, W. & Xiang, A. P. Safeguarding clinical translation of pluripotent stem cells with suicide genes. Organogenesis 9, 34–39 (2013).

24 Tieng, V. et al. Elimination of proliferating cells from CNS grafts using a Ki67 promoter-driven thymidine kinase. Mol Ther Methods Clin Dev 6, 16069 (2016). 10.1038/mtm.2016.69

25 Law, K. C. L., et al. A Selective, Hydrogel-Based Prodrug Delivery System Efficiently Activates a Suicide Gene to Remove Undifferentiated Human Stem Cells Within Neural Grafts. Advanced Functional Materials 33 (2023). 10.1002/adfm.202305771

26 de Luzy, I. R. et al. Human stem cells harboring a suicide gene improve the safety and standardisation of neural transplants in Parkinsonian rats. Nat Commun 12, 3275 (2021). 10.1038/s41467-021-23125-9

27 Liang, Q. et al. Linking a cell-division gene and a suicide gene to define and improve cell therapy safety. Nature 563, 701–704 (2018).

28 Hara, A. et al. Neuron-like differentiation and selective ablation of undifferentiated embryonic stem cells containing suicide gene with Oct-4 promoter. Stem Cells Dev 17, 619–627 (2008). 10.1089/scd.2007.0235

29 Ou, W., Li, P. & Reiser, J. Targeting of herpes simplex virus 1 thymidine kinase gene sequences into the OCT4 locus of human induced pluripotent stem cells. PLoS One 8, e81131 (2013). 10.1371/journal.pone.0081131

30 Yagyu, S., Hoyos, V., Del Bufalo, F. & Brenner, M. K. An Inducible Caspase-9 Suicide Gene to Improve the Safety of Therapy Using Human Induced Pluripotent Stem Cells. Mol Ther 23, 1475–1485 (2015). 10.1038/mt.2015.100

31 Lim, W. A. & June, C. H. The Principles of Engineering Immune Cells to Treat Cancer. Cell 168, 724–740 (2017). 10.1016/j.cell.2017.01.016

32 Khalil, A. S. & Collins, J. J. Synthetic biology: applications come of age. Nat Rev Genet 11, 367–379 (2010). 10.1038/nrg2775

33 Kirkeby, A. et al. Preclinical quality, safety, and efficacy of a human embryonic stem cell-derived product for the treatment of Parkinson’s disease, STEM-PD. Cell Stem Cell 30, 1299–1314 e1299 (2023). 10.1016/j.stem.2023.08.014

34 Pavan, C. et al. A cloaked human stem-cell-derived neural graft capable of functional integration and immune evasion in rodent models. Cell Stem Cell 32, 710–726 e718 (2025). 10.1016/j.stem.2025.03.008

35 de Luzy, I. R. et al. Identifying the optimal developmental age of human pluripotent stem cell-derived midbrain dopaminergic progenitors for transplantation in a rodent model of Parkinson’s disease. Exp Neurol 358, 114219 (2022). 10.1016/j.expneurol.2022.114219

36 Wang, T. et al. snPATHO-seq, a versatile FFPE single-nucleus RNA sequencing method to unlock pathology archives. Commun Biol 7, 1340 (2024). 10.1038/s42003-024-07043-2

37 Song, J. J. et al. Cografting astrocytes improves cell therapeutic outcomes in a Parkinson’s disease model. J Clin Invest 128, 463–482 (2018). 10.1172/JCI93924

38 Yang, F. et al. Activated astrocytes enhance the dopaminergic differentiation of stem cells and promote brain repair through bFGF. Nat Commun 5, 5627 (2014). 10.1038/ncomms6627

39 Serapide, M. F. et al. Boosting Antioxidant Self-defenses by Grafting Astrocytes Rejuvenates the Aged Microenvironment and Mitigates Nigrostriatal Toxicity in Parkinsonian Brain via an Nrf2-Driven Wnt/beta-Catenin Prosurvival Axis. Front Aging Neurosci 12, 24 (2020). 10.3389/fnagi.2020.00024

40 Sozzi, E. et al. Co-grafting strategies uncover cell type–dependent regulation of dopamine neuron specification and functional maturation in a pre-clinical model of Parkinson’s Disease. BioRxiv (2025). 10.64898/2025.12.19.695615

41 Bye, C. R., Rytova, V., Alsanie, W. F., Parish, C. L. & Thompson, L. H. Axonal Growth of Midbrain Dopamine Neurons is Modulated by the Cell Adhesion Molecule ALCAM Through Trans-Heterophilic Interactions with L1cam, Chl1, and Semaphorins. J Neurosci 39, 6656-6667 (2019). 10.1523/JNEUROSCI.0278-19.2019

42 Eyquem, J. et al. Targeting a CAR to the TRAC locus with CRISPR/Cas9 enhances tumour rejection. Nature 543, 113–117 (2017). 10.1038/nature21405

43 Tiklova, K. et al. Single cell transcriptomics identifies stem cell-derived graft composition in a model of Parkinson’s disease. Nat Commun 11, 2434 (2020). 10.1038/s41467-020-16225-5

44 Storm, P. et al. Lineage tracing of stem cell-derived dopamine grafts in a Parkinson’s model reveals shared origin of all graft-derived cells. Sci Adv 10, eadn3057 (2024). 10.1126/sciadv.adn3057

45 Kotini, A. G. et al. Stage-Specific Human Induced Pluripotent Stem Cells Map the Progression of Myeloid Transformation to Transplantable Leukemia. Cell Stem Cell 20, 315–328 e317 (2017). 10.1016/j.stem.2017.01.009

46 Pang, Z. P. et al. Induction of human neuronal cells by defined transcription factors. Nature 476, 220–223 (2011). 10.1038/nature10202

47 Lu, C. & Sanjana, N. E. Generation of a knock-in MAP2-tdTomato reporter human embryonic stem cell line with inducible expression of NEUROG2/1 (NYGCe001-A). Stem Cell Res 41, 101643 (2019). 10.1016/j.scr.2019.101643

48 Boutin, C. et al. NeuroD1 induces terminal neuronal differentiation in olfactory neurogenesis. Proc Natl Acad Sci U S A 107, 1201–1206 (2010). 10.1073/pnas.0909015107

49 Khan, S. et al. Survival of a Novel Subset of Midbrain Dopaminergic Neurons Projecting to the Lateral Septum Is Dependent on NeuroD Proteins. J Neurosci 37, 2305–2316 (2017). 10.1523/JNEUROSCI.2414-16.2016

50 Clackson, T. et al. Redesigning an FKBP-ligand interface to generate chemical dimerizers with novel specificity. Proc Natl Acad Sci U S A 95, 10437–10442 (1998). 10.1073/pnas.95.18.10437

51 Gantner, C. W., Cota-Coronado, A., Thompson, L. H. & Parish, C. L. An Optimized Protocol for the Generation of Midbrain Dopamine Neurons under Defined Conditions. STAR Protocols 1, 100065 (2020). 10.1016/j.xpro.2020.100065

52 Moriarty, N. et al. Understanding the Influence of Target Acquisition on Survival, Integration, and Phenotypic Maturation of Dopamine Neurons within Stem Cell-Derived Neural Grafts in a Parkinson’s Disease Model. J Neurosci 42, 4995–5006 (2022). 10.1523/JNEUROSCI.2431-21.2022

53 Hao, Y. et al. Dictionary learning for integrative, multimodal and scalable single-cell analysis. Nat Biotechnol 42, 293–304 (2024). 10.1038/s41587-023-01767-y

54 Becht, E. et al. Dimensionality reduction for visualizing single-cell data using UMAP. Nat Biotechnol (2018). 10.1038/nbt.4314

55 Stuart, T. et al. Comprehensive Integration of Single-Cell Data. Cell 177, 1888–1902 e1821 (2019). 10.1016/j.cell.2019.05.031

56 Ianevski, A., Giri, A. K. & Aittokallio, T. Fully-automated and ultra-fast cell-type identification using specific marker combinations from single-cell transcriptomic data. Nat Commun 13, 1246 (2022). 10.1038/s41467-022-28803-w

57 Yu, G., Wang, L. G., Han, Y. & He, Q. Y. clusterProfiler: an R package for comparing biological themes among gene clusters. OMICS 16, 284–287 (2012). 10.1089/omi.2011.0118

58. Gentleman, R. C., et al. Bioconductor: open software development for computational biology and bioinformatics. Genome Biol 5, R80 (2004). 10.1186/gb-2004-5-10-r80

59 Ashburner, M. et al. Gene ontology: tool for the unification of biology. The Gene Ontology Consortium. Nat Genet 25, 25–29 (2000). 10.1038/75556

60 Gu, Z., Eils, R. & Schlesner, M. Complex heatmaps reveal patterns and correlations in multidimensional genomic data. Bioinformatics 32, 2847–2849 (2016). 10.1093/bioinformatics/btw313

